# MinJ is a conserved nine-pass transmembrane protein that contains a putative transmembrane β-sheet

**DOI:** 10.64898/2026.07.02.736092

**Authors:** Kehinde O. Adebiyi, Laura C. Lastra, Jane H. Joncha, Sadie B. Ruesewald, Stephen C. Jacobson, Daniel B. Kearns

**Affiliations:** Department of Biology, Indiana University, Bloomington IN USA; Department of Chemistry, Indiana University, Bloomington IN USA

## Abstract

The Min system disassembles FtsZ-rings after septation in *Bacillus subtilis* and is localized to the nascent division plane and cell poles by the protein MinJ. The N-terminal region of MinJ contains transmembrane segments while the C-terminal region of MinJ contains a PDZ domain but its topology and functional domains are poorly understood. Here we empirically test MinJ topology based on a variety of transmembrane prediction models and find that the data is most consistent with Alphafold3, which predicts a 9-pass transmembrane protein with an external N-terminus and internal C-terminus. Deletion analysis indicates that all regions of the protein tested are required for function but deletion of the PDZ domain alone preserves polar localization and interaction with both MinD and DivIVA. Moreover, Alphafold predicts that transmembrane segments 6 and 7 comprise staves of an unusual transmembrane β-sheet and deletion of the putative β-sheet in the absence of MinD results in a minicell frequency that exceeds mutation of MinD alone. Bioinformatic analysis indicates that MinJ is highly conserved within Firmicutes and is co-conserved with MinD and DivIVA with which it interacts. Our data clarify the structure of MinJ and support models in which MinJ has functions in addition to restricting the activity of the Min system.

**IMPORTANCE:** Faithful positioning of the bacterial division site is important for cell growth and is coordinated by the conserved Min system. Although the Min system of *Bacillus subtilis* has been extensively studied, MinJ, the membrane protein that links the division inhibitor MinCD to the polar determinant DivIVA, remains the least well-understood. Here we experimentally define the membrane topology of MinJ and show that our data are most consistent with a nine-pass transmembrane architecture predicted by AlphaFold3. We further provide genetic, cell biological, and evolutionary evidence supporting that two of the staves form a highly conserved putative transmembrane β-sheet, a structure normally excluded from the plasma membrane. Our findings refine MinJ structural organization and provide a framework for understanding its conserved functions in bacterial cell division.

## INTRODUCTION

Bacterial cell division is initiated by the polymerization of the tubulin homolog FtsZ into protofilaments that dynamically treadmill and coalesce into a structure called the Z-ring (1–5). The Z-ring intensifies over time and once mature, it recruits transmembrane divisome proteins that drive peptidoglycan synthesis and membrane invagination during septation (6–10). In the Gram-positive model organism *Bacillus subtilis*, a membrane-associated complex of MinC and MinD disassembles the Z-ring after septation to recycle FtsZ monomers for the next round of division (11–14). The MinCD complex is localized to the nascent division plane and retained at the cell poles by the action of a third protein, DivIVA, a tropomyosin-like coiled-coil protein that oligomerizes and is recruited to membrane regions with increased negative curvature (15–19). No direct interaction between DivIVA and either MinC or MinD was detected however, and a fourth protein, MinJ, was proposed to act as an intermediary (20–21).

Multiple lines of evidence support the conclusion that MinJ acts as a linker to connect the MinCD cell division inhibitor to the DivIVA polar localization complex. First, mutation of MinJ produced filamentous cells due to a defect in cell division, phenocopying the absence of DivIVA, and cell division could be restored by mutation of MinCD (15; 20-22). Second, MinJ localized to nascent division sites and cell poles in a DivIVA-dependent manner, and MinCD localization was dependent on MinJ (20–21). Finally, pairwise combinations of proteins expressed in *Escherichia coli* as part of a bacterial two-hybrid system indicated positive interactions for MinJ when paired with itself, MinD and DivIVA (20–21). Thus, MinJ appeared to act as a structural linker between the two systems. Early bioinformatic analysis indicated that the MinJ N-terminal region contained multiple transmembrane segments and that the C-terminal region contained PDZ domain (20–21). Precisely how MinJ restricts the localization and activity of MinCD to the cell pole remains poorly understood.

Subsequent work in *B. subtilis* suggested a more complex role for MinJ. For example, one study suggests that not only does MinJ control Z-ring stability through MinCD, but may also play a second role in the disassembly of the divisome after septation (23). Moreover, protein pulldowns using the MinJ soluble PDZ domain did not pull down either MinD or DivIVA, but purified a number of proteins relevant to cell division instead, perhaps pointing to a second parallel function (24). Yet another study suggested that MinJ directly interacted with RodZ, a protein involved in cellular elongation seemingly unrelated to cell division (25–27). Complicating matters further, while *Clostridium beijerinckii* and *Clostridium difficile* encode MinJ, direct interaction was detected between MinD and DivIVA, seemingly obviating the need of a linker protein (28). Moreover, MinJ in *Listeria monocytogenes* localized to the cell poles in a DivIVA-independent manner, but mutation of MinJ did not give rise to a cell division defect (29). Thus, the function of MinJ may be more complex than previously realized, and vary across the phylogenetic tree.

To further explore the function in *B. subtilis*, we sought to determine the topology of MinJ in the membrane and define its functional domains. Using a combination of genetic, cell biological, and bioinformatic approaches, we demonstrate that MinJ is a nine-pass transmembrane protein with a cytoplasmic PDZ domain. Moreover, topological analysis was consistent with a structure predicted by Alphafold3 in which two of the nine transmembrane segments form a β-sheet. Deletion of various parts of the protein resulted in a loss-of-function filamentous phenotype, but deletion of the putative transmembrane β-sheet gave rise to a hyper minicell frequency that was additive when MinD was simultaneously mutated. Deletion of the PDZ domain also abolished MinJ function but maintained both subcellular localization and interaction with both MinD and DivIVA; thus if the PDZ is an interaction domain, its interaction is with another binding partner. Finally, bioinformatic analysis supports high co-conservation of MinJ, MinD, and DivIVA in the Firmicutes, and Alphafold3 structural comparisons supports a high degree of structural conservation. Our work functionally supports the role of MinJ in regulating cell division site selection and points to additional functions that may perhaps reconcile discrepancies found in other related organisms.

## RESULTS

### MinJ is a nine-pass transmembrane protein with a cytoplasmic PDZ domain

MinJ appears to be a multi-pass transmembrane protein with a C-terminal PDZ domain but the number of transmembrane segments it contains, and its overall topology is unknown. Transmembrane segments were predicted with the bioinformatic sequence analysis programs TM-PRED, UNIPROT, DeepHMTMM (**Fig 1A, Fig S1**), and the structural program Alphafold3 **(Fig 1B**) which predicted that MinJ had 6, 7, 9, and 9 transmembrane segments, respectively. To empirically determine both the number of transmembrane segments as well as membrane topology, N-terminal MinJ fragments were expressed from the IPTG-inducible *P_hyspank_*promoter, translationally fused to either the *lacZ* gene encoding β-galactosidase or the *phoA* gene encoding alkaline phosphatase, and inserted at an ectopic site in the chromosome (30). As LacZ is only active in the cytoplasm and PhoA is only active extracellularly, the strains should give opposing activity based on the topology of the transmembrane segments immediately preceding the fusion (31).

**Figure 1.**
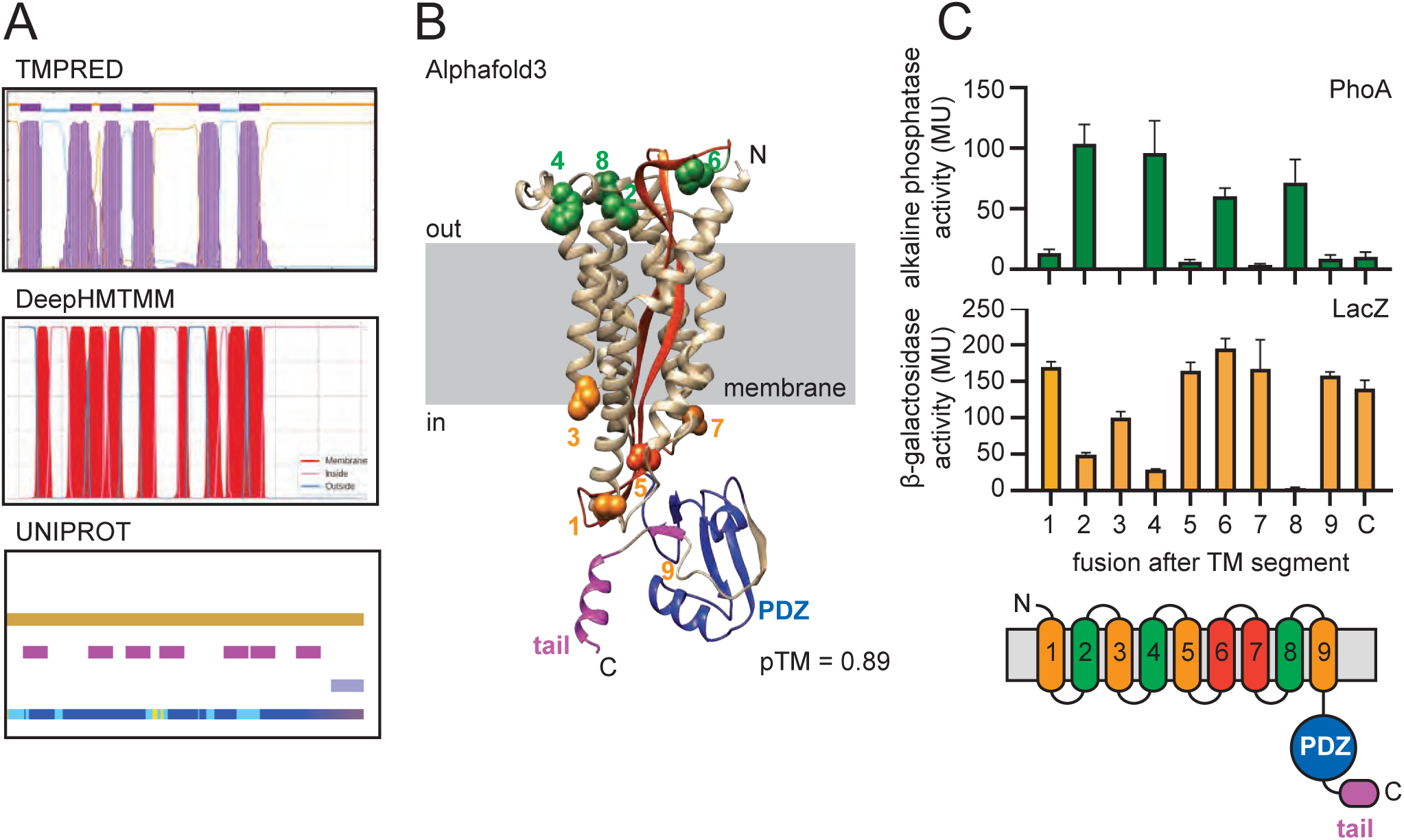
MinJ is a nine-pass transmembrane protein with a cytoplasmic PDZ domain. **A)** Topology predictions of MinJ generated using the transmembrane prediction programs TMPRED, DeepTMHMM, and UniProt. **B)** AlphaFold3 structural model of *Bacillus subtilis* MinJ overlaid on a cartoon representation of the plasma membrane. The predicted transmembrane domain consists of seven α-helical transmembrane segments shown in gold and two β-sheet transmembrane staves shown in red. The C-terminal PDZ domain is shown in blue, and the extended C-terminal tail is shown in purple. Green spheres indicate positions at which translational fusions to PhoA gave positive activity and orange spheres indicate positions at which translational fusions to LacZ gave positive activity, shown in panel C. **C)** Enzymatic activity measurement of PhoA (top, green bars) and LacZ (bottom, orange bars) of fusions made after the indicated transmembrane segments predicted by Alphafold3 (cartoon). Each bar represents the average Miller units of three biological replicates and error bars indicate standard deviation. The following strains were used to generate these data: PhoA fusions; TM1 (DK9628), TM2 (DK9872), TM3 (DK9627), TM4 (DK9626), TM5 (DK9625), TM6 (DK9670), TM7 (DK9624), TM8 (DK9671), TM9 (DK9623) and PDZ (DK9874). LacZ fusions; TM1 (DK9583), TM2 (DK9866), TM3 (DK9582), TM4 (DK9581), TM5 (DK9580), TM6 (DK9668), TM7 (DK9494), TM8 (DK9669), TM9 (DK9495) and PDZ (DK9791).

Quantitative enzymatic assays for β-galactosidase (LacZ) and alkaline phosphatase (PhoA) were measured in the presence of their cognate chromogenic substrates ONPG and ONPP, respectively. Each of the programs predicted the first transmembrane segment similarly, and fusions after the first putative transmembrane segment indicated a cytoplasmic facing (**Fig 1A,B**) placing the MinJ N-terminus in the extracellular space (**Fig S1, Fig 1C**). Empirical testing of each bioinformatic model with fusions made after subsequent transmembrane segments however, failed to give alternating activities that one might expect for sequential transmembrane passes (**Fig S1**). The data best fit the Alphafold3 model with only one anomalous result where both LacZ and PhoA appeared to be simultaneously active when inserted after putative transmembrane segment 6 (**Fig 1C**). We infer that the end of transmembrane segment 6 is extracytoplasmic as it is flanked by the termini of transmembrane segments 5 and 7 that both support cytoplasmic localization (**Fig 1C**). Moreover, PhoA can only be active if the fusion is in the extracellular space whereas LacZ can give a false positives due to folding defects. We note that the separation between transmembrane segment 6 and 7 consists of only a single proline and either the tight turn resulted in stochastic insertion of the anomalous fusion or it was consequence of the architecture predicting that segments 6 and 7 form an unusual transmembrane β-sheet. Finally, empirical analysis indicated that MinJ C-terminal PDZ domain was cytoplasmic, consistent with an extracellular N-terminus and an odd number of transmembrane segments (**Fig S1, Fig 1C**). We conclude that MinJ is a nine-pass transmembrane protein, with a potentially unusual architecture as two of the transmembrane segments may consist of β-sheets.

### The MinJ putative transmembrane β-sheet and PDZ domain are differentially required for function

We next sought to characterize the requirement of various domains for MinJ based on the topological model supported by AlphaFold3. To do so, a complementation construct was generated in which the *minJ* gene was expressed from its native promoter (*P_minJ_*) and inserted at an ectopic site in the chromosome of a *minJ* mutant. Next, sequential pairs of transmembrane segments were deleted from the MinJ complementation construct to maintain the overall topology of the protein with respect to the N- and C- termini, and each was separately integrated in a *minJ* mutant background. Specifically, MinJ allelic variants were generated that deleted helices two and three (MinJ^Δ23^), helices four and five (MinJ^Δ45^), sheet staves six and seven (MinJ ^Δ67^), and lastly, helices eight and nine (MinJ ^Δ89^). In addition, mutants were generated that deleted the C-terminal PDZ (MinJ^ΔPDZ^) and the entire cytoplasmic domain (MinJ^ΔC^). Whereas complementation with the wild type allele of *minJ* restored swarming motility (**Fig S2**) and wild type cell length to the *minJ* mutant, each mutant allele failed to complement either phenotype (**Fig 2, Fig S3**). We conclude that each segment that was deleted was required for MinJ functionality.

**Figure 2.**
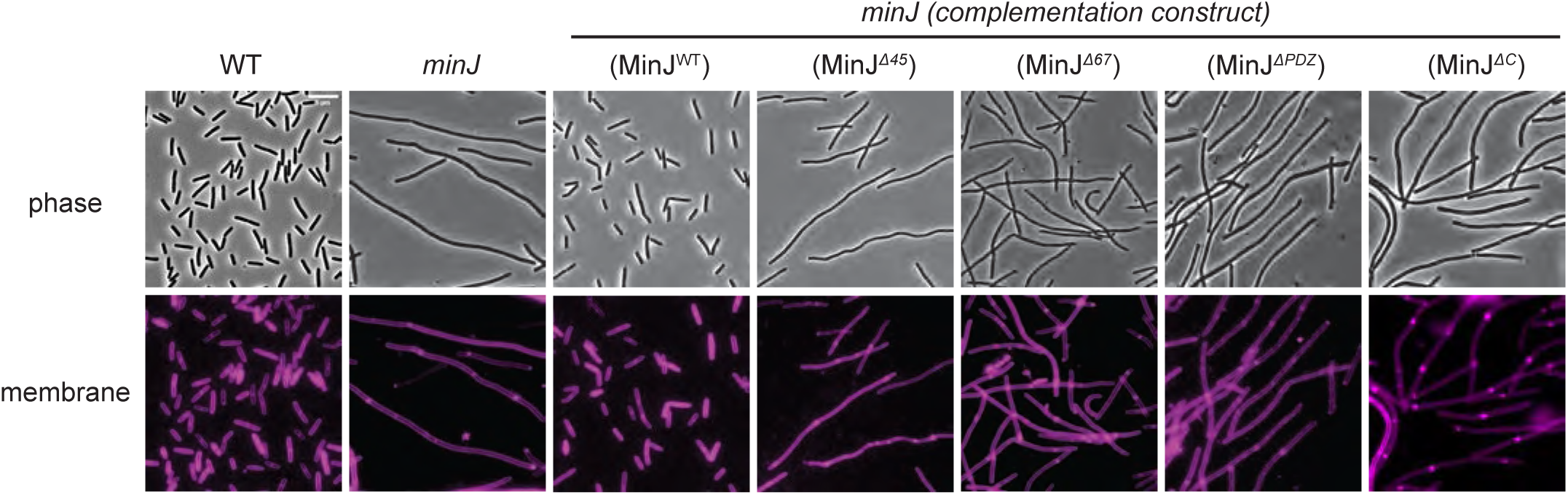
Deletion of pairs of MinJ transmembrane domains and deletion of the C-terminal domain results in a cell division defect. Micrograph images of phase-contrast and fluorescence microscopy of cell membranes stained with FM4-64 (false colored magenta) for the genetic backgrounds indicated. Genetic complementation construct provided *in trans* is indicated in parentheses. The following strains were used to generate these data: WT (DK1042), *minJ* (DB92), *minJ (P_minJ_-minJ)* (DB2484), *minJ (P_minJ_-minJ^Δ45^)* (DB2500), *minJ (P_minJ_- minJ^Δ67^)* (DB3144), *minJ (P_minJ_-minJ^ΔPDZ^)* (DB2502), and *minJ (P_minJ_-minJ^ΔC^)* (DB2485). Scale bar is 5 μm and is the same for all panels.

The various alleles of MinJ could fail to complement a *minJ* null mutation either because they are defective in MinJ activity or because they are defective in MinJ protein stability. To determine which, if any, of the alleles were unstable, protein lysates were collected from each complementation strain, resolved by SDS-PAGE, and subjected to Western blot analysis. Each lysate was probed with an antibody raised against the soluble C-terminal domain of MinJ and separately with an antibody raised against the vegetative sigma factor SigA to serve as a loading control. MinJ protein was present in the wild type, absent in the *minJ* mutant, and restored to wild type levels in the presence of the complementation construct expressing the wild type allele (**Fig 3**). Deletion of the various transmembrane segments decreased the size of the MinJ reactive band. Moreover, the MinJ^Δ23^, MinJ^Δ45^, and MinJ^Δ89^ alleles substantially reduced MinJ protein levels, while the MinJ^Δ67^ allele did not. Finally, the MinJ^ΔPDZ^ and MinJ^ΔC^ alleles were both undetectable and uninterpretable, as the C-terminus to which the antibody was raised was either absent or severely truncated in both (**Fig 3**). We continued with MinJ^Δ45^ and MinJ^Δ67^ alleles as representatives of unstable and stable transmembrane defective alleles respectively, as well as the nested C-terminal deletions for further analysis.

**Figure 3.**
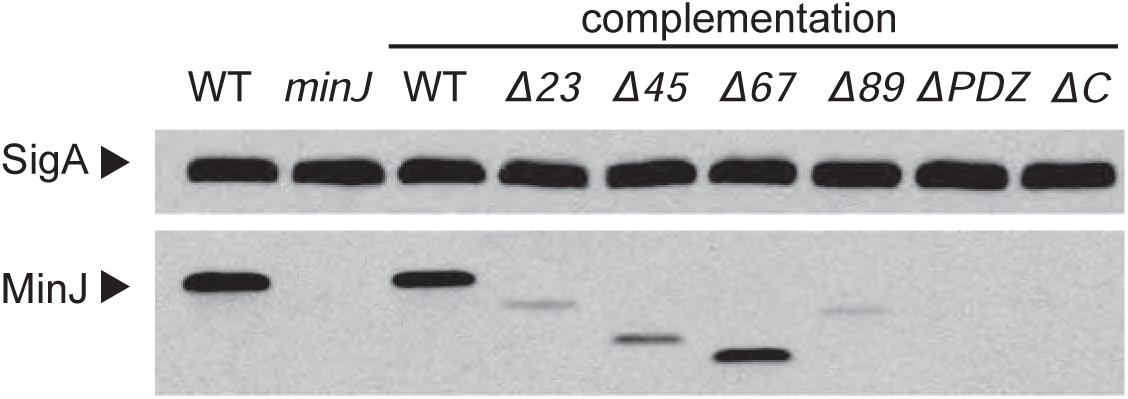
Deletion of transmembrane segments differentially affects MinJ protein accumulation. Western blot analysis of cell lysates isolated from varying *minJ* deletion. Samples were separated by SDS-PAGE and probed with a primary antibody raised against the soluble PDZ domain of MinJ. A primary antibody raised against the vegetative sigma factor SigA was used as a loading control. The following strains were used to generate these data: WT (DK1042), *minJ* (DB92), *minJ (P_minJ_-minJ)* (DB2484), *minJ (P_minJ_-minJ^Δ23^)* (DB2499), minJ *(P_minJ_-minJ^Δ45^)* (DB2500), *minJ (P_minJ_-minJ^Δ67^)* (DB3144), *minJ (P_minJ_-minJ^Δ89^)* (DB2501), *minJ (P_minJ_-minJ^ΔPDZ^)* (DB2502), and *minJ (P_minJ_-minJ^ΔC^)* (DB2485).

MinJ localizes to nascent division septa and cell poles (Patrick 2008; Bramkamp 2008). To further explore the domains of MinJ, localization of mutated alleles was tested by fusing the fluorescent protein mNeongreen to the C-terminus of the complementation construct and integrated at an ectopic site. Furthermore, the native gene encoding MinJ was deleted in each strain so as not to compete with the fusion for localization, and MinD was mutated to increase the frequency of division events and cell poles to aid interpretation. Whereas wild type MinJ localized to cell poles, fusion of mNeongreen to MinJ^Δ45^, MinJ^Δ67^, and deletion of the entire MinJ^ΔC^ C-terminal domain abolished localization such that fluorescence appeared as a diffuse haze (**Fig 4**). Mutation of the MinJ PDZ domain alone, however, localized more like the wild type with clear polar enrichment (**Fig 4**). We conclude that localization of MinJ requires the transmembrane region and the entire C-terminus but not the PDZ domain specifically.

**Figure 4.**
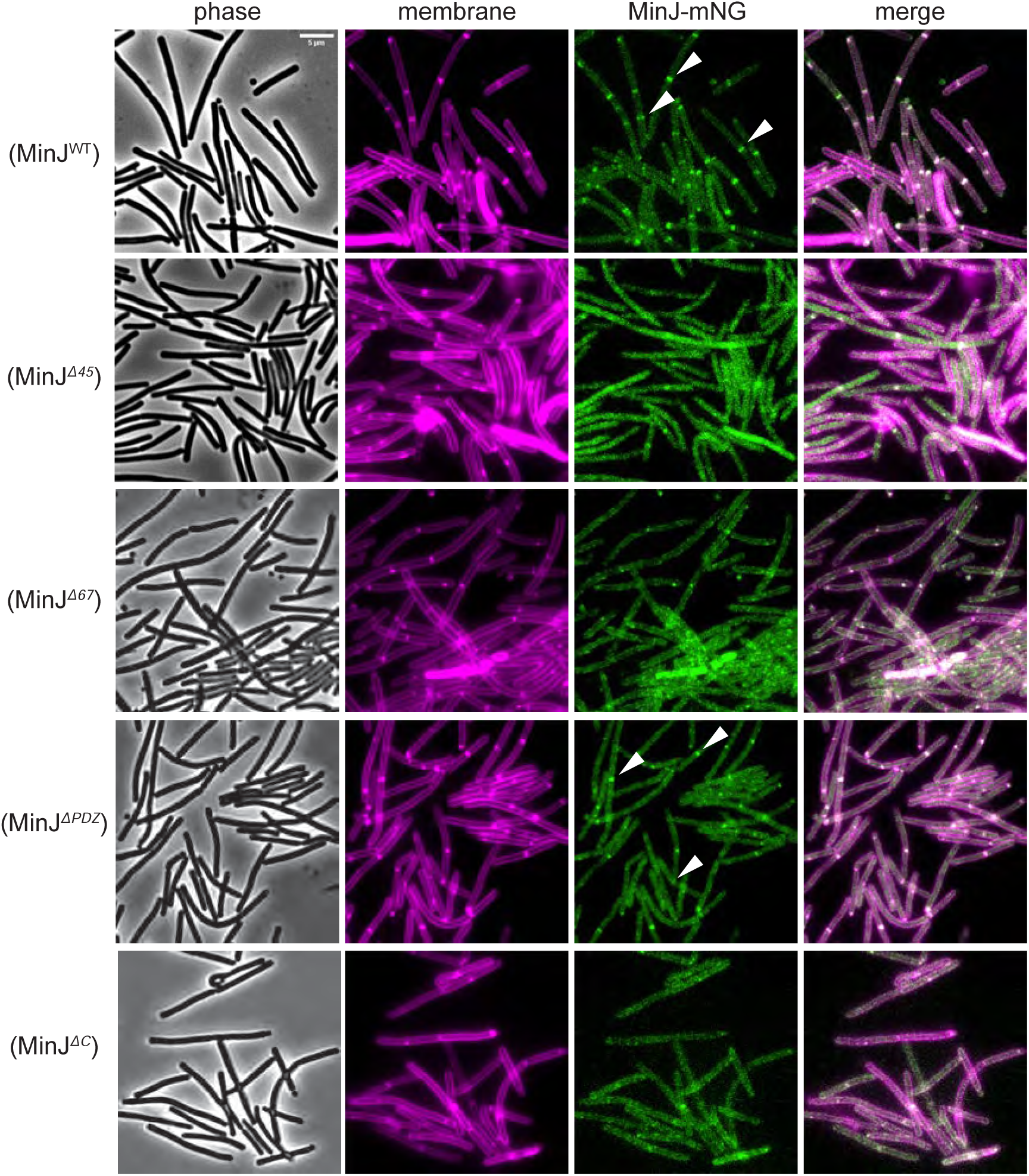
Localization of MinJ requires the transmembrane domain and the C-terminal region but not the PDZ domain, specifically. Micrograph images of phase-contrast microscopy, fluorescence microscopy of cell membranes stained with FM4-64 (false colored magenta), MinJ-mNeongreen localization (false colored green), and a merge of the two fluorescence channels for the genetic backgrounds indicated. All MinJ-mNeongreen fusions were expressed from the native *P_minJ_* promoter at an ectopic chromosomal locus in a *minJ* mutant background. In addition, each strain was mutated for MinD to increase the frequency of division septa and cell poles and thereby facilitate localization analysis. Carets indicate polar localization events. The following strains were used to generate these data: *minD minJ (P_minJ_-minJ-mNeongreen)* (DB3506), *minD minJ (P_minJ_-minJ^Δ45^-mNeongreen)* (DB3582), *minD minJ (P_minJ_-minJ^Δ67^-mNeongreen)* (DB3508*), minD minJ (P_minJ_-minJ^ΔPDZ^-mNeongreen)* (DB3509), and *minD minJ (P_minJ_-minJ^ΔC^-mNeongreen)* (DB3626). Scale bar is 5 μm and is the same for all panels.

MinJ is thought to localize to the cell pole by direct interaction with the polar determinant DivIVA, and in turn, control cell division site selection by interacting with, and sequestering, MinD. To determine whether the loss-of-function alleles disrupted multimerization and/or interaction DivIVA and MinD, each allele was fused to a fragment of adenylate cyclase and expressed in *E. coli* with a complementary fragment fused to an interacting partner. In this bacterial two-hybrid assay, a positive interaction between the two proteins restores adenylate cyclase activity which in turn activates β-galactosidase gene expression. Consistent with previous reports, wild type MinJ interacted with itself, MinD and DivIVA, as β-galactosidase activity of the pairwise combinations dramatically exceeded controls in which nothing was fused to each half of adenylate cyclase (**Fig 5A**). Deletion of the MinJ transmembrane domains 4 and 5 (MinD^Δ45^), and 6 and 7 (MinD^Δ67^) abolished interaction with each protein tested (**Fig 5B,C**). Mutation of the PDZ domain abolished interaction with MinJ, but maintained partial interaction with both DivIVA and MinD, perhaps consistent with the fact that polar localization was observed with this allele (**Fig 5D**). Finally, deletion of the entire C-terminus exhibited an interaction pattern similar to that of the PDZ deletion, albeit with stronger levels of expression (**Fig 5E**). We conclude that the transmembrane domain is both necessary and sufficient for interaction with DivIVA and MinD, at least in the heterologous bacterial two-hybrid system.

**Figure 5.**
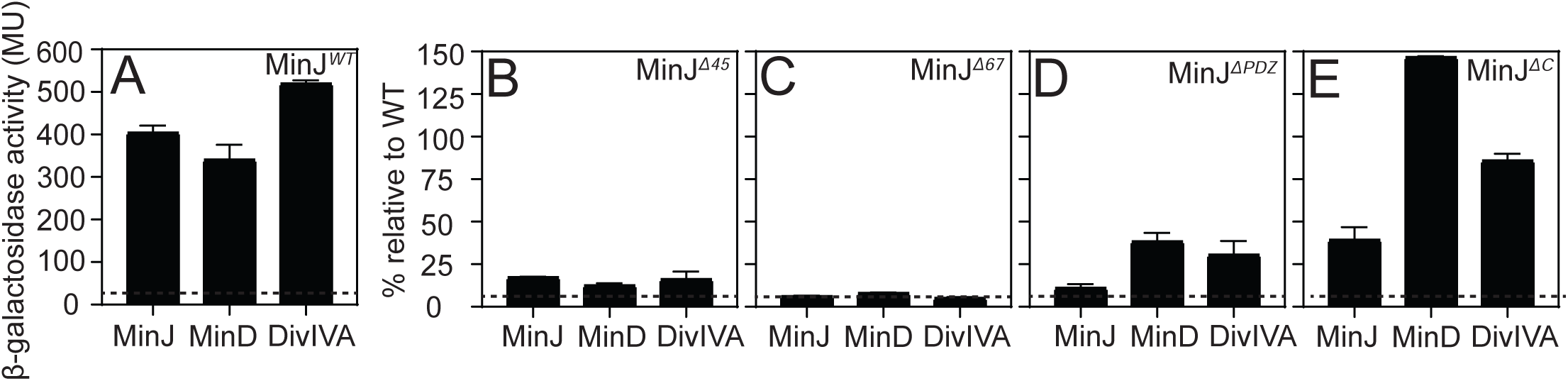
The MinJ transmembrane domain, but not the C-terminal domain, is required for interaction with MinD, DivIVA, and MinJ. Protein-protein interactions were measured using an *E. coli*-based bacterial two-hybrid assay. In this assay, proteins of interest were fused to complementary fragments of the *Bordetella pertussis* adenylate cyclase enzyme. Interaction between the fusion proteins restores adenylate cyclase activity, resulting in expression of β-galactosidase. **A)** β-galactosidase activity, expressed as Miller units, for pairwise interactions between wildtype MinJ and the proteins indicated below the x-axis. Wild type MinJ interacted with itself, MinD, and DivIVA, whereas controls lacking a fusion partner exhibited only background activity. The dotted horizontal line indicates the activity of the empty-vector control. **(B–E)** Interaction strength of MinJ mutant alleles MinJ^Δ45^ (panel B), MinJ^Δ67^ (panel C), MinJ^ΔPDZ^ (panel D), and MinJ^ΔC^ (panel E), expressed as a percentage of the corresponding wildtype interaction shown in Panel A. Percent interaction was calculated by dividing the Miller units obtained for a given mutant interaction by the mean Miller units obtained for the corresponding wildtype interaction and multiplying by 100. The dotted horizontal line indicates the activity of the empty-vector control expressed relative to the corresponding wildtype interaction. Each bar represents the average of three biological replicates and error bars indicate standard deviation. The following *E. coli* strains were used to generate these data: **A)** *minJ-CyaAT18/minJ-CyaAT25* (DE4545), *minJ-CyaAT18/minD-CyaAT25* (DE4552), *minJ-CyaAT18/divIVA-CyaAT25* (DE4559), and the empty vector control *CyaAT18/CyaAT25* (DE4566). B) *minJ^Δ45^ -CyaAT18/minJ^Δ45^-CyaAT25* (DE4546), *minJ^Δ45^-CyaAT18/minD-CyaAT25* (DE4553), and *minJ^Δ45^-CyaAT18/divIVA-CyaAT25* (DE4560). C) *minJ^Δ67^-CyaAT18/minJ^Δ67^-CyaAT25* (DE4547), *minJ^Δ67^-CyaAT18/minD-CyaAT25* (DE4554), and *minJ^Δ67^-CyaAT18/divIVA-CyaAT25* (DE4561). D) *minJ^ΔPDZ^-CyaAT18/minJ^ΔPDZ^-CyaAT25* (DE4548), *minJ^ΔPDZ^-CyaAT18/minD-CyaAT25* (DE4555), and *minJ^ΔPDZ^-CyaAT18/divIVA-CyaAT25* (DE4562). E) *minJ^ΔC^-CyaAT18/minJ^ΔC^-CyaAT25* (DE4550), *minJ^ΔC^-CyaAT18/minD-CyaAT25* (DE4557), and *minJ^ΔC^-CyaAT18/divIVA-CyaAT25* (DE4564).

### MinJ sequence conservation and structural diversity

To contextualize the topological and functional analyses, we next examined the MinJ phylogenetic distribution and sequence conservation. To assess sequence conservation, a phylogenetic tree spanning species from all domains of life was generated with PhyloT, with Bacteria represented broadly across major phyla. We then systematically queried the NCBI Protein database for the presence of MinJ in each species using its taxonomic identifier and found that MinJ homologs were detected exclusively within the Firmicutes (“Bacillota”) **(Fig. 6A)**. In parallel, we assessed structural conservation, where the AlphaFold3-predicted structure of *Bacillus subtilis* MinJ input to Foldseek identified 2,480 structurally related proteins within Bacillota and only four distant clusters within Actinomycetota **(Fig. S4)**. The latter were sparse and did not form a coherent clade, suggesting either remote structural similarity or potential false positives rather than true orthology. We conclude that MinJ is highly conserved within Firmicutes and largely absent from other bacterial phyla.

**Figure 6.**
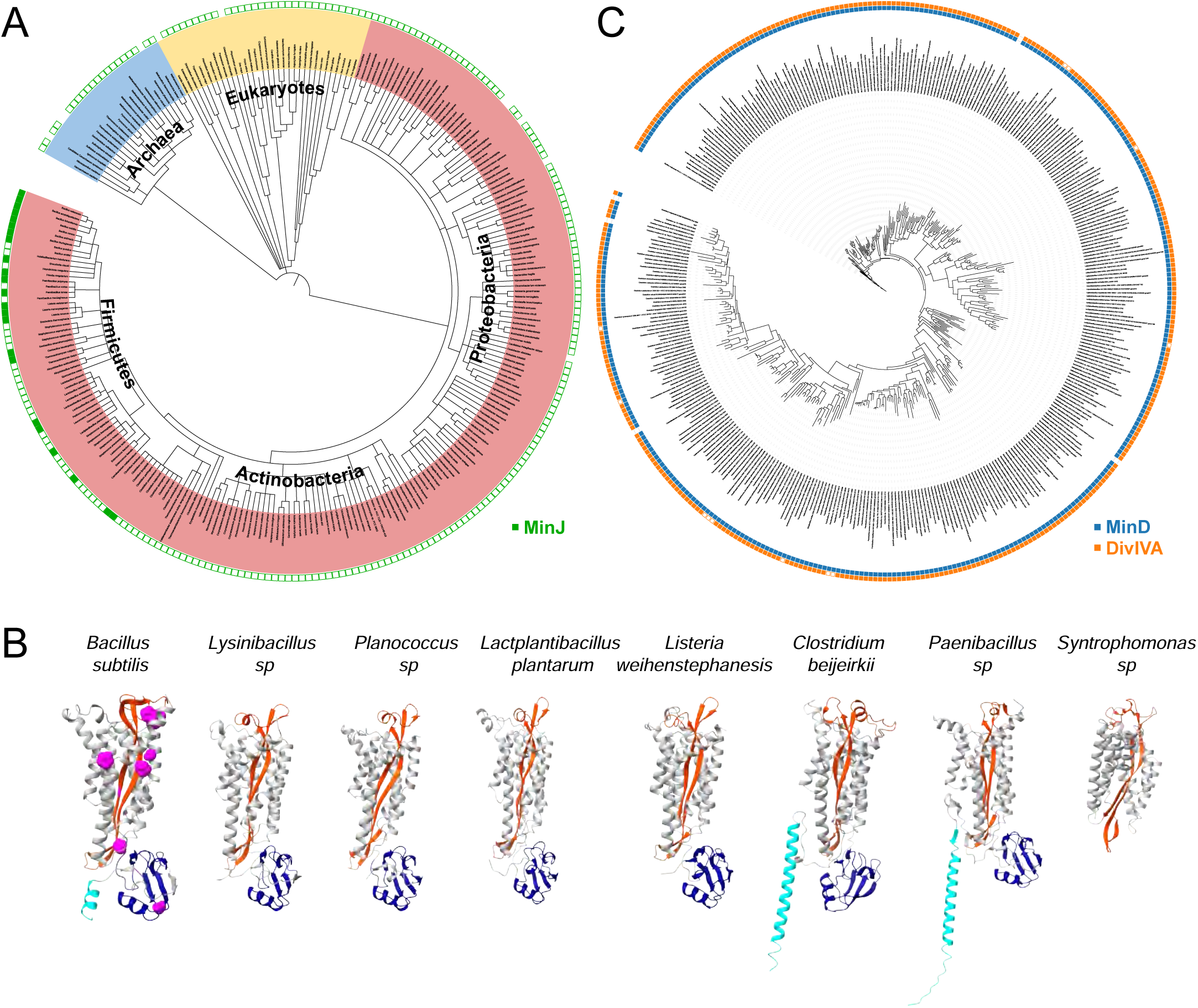
MinJ is restricted to the Bacillota, co-occurs with MinD and DivIVA, and is predicted to have high structural conservation. **A)** Phylogenetic tree spanning representative species from Bacteria, Archaea, and Eukarya. Archaea are highlighted in light blue, Eukarya in yellow, and Bacteria in pink. The presence or absence of MinJ was determined by querying the NCBI Protein database using species-specific taxonomic identifiers. Filled green squares indicate the presence of a MinJ homolog, whereas open squares indicate the absence of a MinJ homolog. **B)** AlphaFold3 structural models of representative MinJ orthologs selected from progressively more distant branches of the MinJ phylogeny. Structures are shown for *Bacillus subtilis*, *Lysinibacillus*, *Planococcus*, *Lactiplantibacillus*, *Listeria*, *Clostridium*, *Paenibacillus*, and *Syntrophomonas*. Predicted α-helical transmembrane segments are shown in light gray, β-sheet transmembrane staves are shown in orange, the PDZ domain is shown in blue, and the extended C-terminal tail is shown in cyan. Residues exhibiting greater than 90% conservation across the 396 MinJ orthologs are mapped onto the *Bacillus subtilis* MinJ structure and represented as magenta spheres. **C)** Maximum-likelihood phylogenetic tree of 396 MinJ orthologs identified using EggNOG and reconstructed using IQ-TREE. Blue annotation tracks indicate the presence or absence of MinD and orange annotation tracks indicate the presence or absence of DivIVA. Filled squares represent gene presence and open squares represent gene absence.

To further delineate conserved features, 396 MinJ orthologs were used to generate a multiple sequence alignment and a model-based phylogenetic reconstruction. Eleven individual residues distributed throughout the protein exhibited greater than 90% conservation **(Fig 6B, magenta; Fig S5)**. One of the residues was a proline preceded by two tryptophans (WWP) and the trio were conserved in 73% of sequences **(Table S1)**. Structural mapping of the WWP motif onto the AlphaFold3 model of *B. subtilis* MinJ places this sequence within the turn separating the putative β-sheet transmembrane segments 6 and 7, corresponding to the unusual β-sheet architecture identified in our topological analysis **(Fig S5).** The strong conservation of this tryptophan-rich proline-turn suggests that the putative transmembrane β-sheet is evolutionarily distributed, and structural comparison of seven phylogenetically diverse MinJ representatives supported a conserved transmembrane architecture (**Fig 6B**).

Finally, as MinJ is thought to function in concert with the cell division proteins MinD and DivIVA, we assessed the co-occurrence of MinD and DivIVA in species encoding MinJ. Querying the NCBI protein database for each orthologous species revealed that nearly all species that encode MinJ also encode MinD and DivIVA **(Fig 6C).** Some rare species that encode MinJ and MinD lack a DivIVA homolog, but in these cases, AlphaFold3-predicted structures of the MinJ orthologs with a helical extended C-terminal extension resembling the coiled-coil domain of DivIVA (**Fig 6B, cyan**). Perhaps in these cases, the two proteins, MinJ and DivIVA, may have been fused into one. Finally, rare cases were observed in which MinJ lacked a C-terminal PDZ domain, but if or how the lack of the PDZ was compensated for was unclear. Taken together, we conclude that MinJ is a Firmicute-restricted protein with a highly conserved membrane topology and domain architecture, and that its evolutionary trajectory is closely linked to the Min system components with which it functionally interacts.

### Deletion of the MinJ putative transmembrane β-sheet confers a hyper-minicell phenotype

During microscopy analysis of the mutant alleles of MinJ, we noticed that deletion of transmembrane segments 6 and 7 produced minicells and minicells appeared unusually abundant when MinD was also disrupted. To quantify the minicell frequency, cells separately disrupted for MinD, MinJ, and simultaneously disrupted for both were grown in microfluidic channels and the number of divisions giving rise to minicells were divided by the total number of division events observed. Cells lacking MinD exhibited a high frequency of minicells, cells lacking MinJ exhibited a low frequency of minicells, and mutation of *minD* was epistatic such that the *minD minJ* double mutant resembled the absence of MinD alone **(Fig 7A)**. Next, each mutant allele of MinJ was introduced at an ectopic site of the *minD minJ* double mutant background such that it was the only copy of MinJ in the cell. Cells expressing MinJ^Δ45^, MinJ^ΔPDZ^, and MinJ^ΔC^, all phenocopied deletion of MinD, but cells expressing MinJ^Δ67^ exhibited a hyperminicell phenotype that was statistically greater than mutation of *minD* (**Fig 7A**). Moreover, the minicells of the *minD* MinJ^Δ67^ double mutant were larger than the minicells produced when MinD was absent alone (**Fig 7B**). We conclude that transmembrane segments 6 and 7 play an important role in MinJ function and likely contribute to a second mechanism by which MinJ suppresses cell division. Thus, the putative transmembrane β-sheet is not only a highly conserved structural element but plays a central role governing MinJ activity.

**Figure 7.**
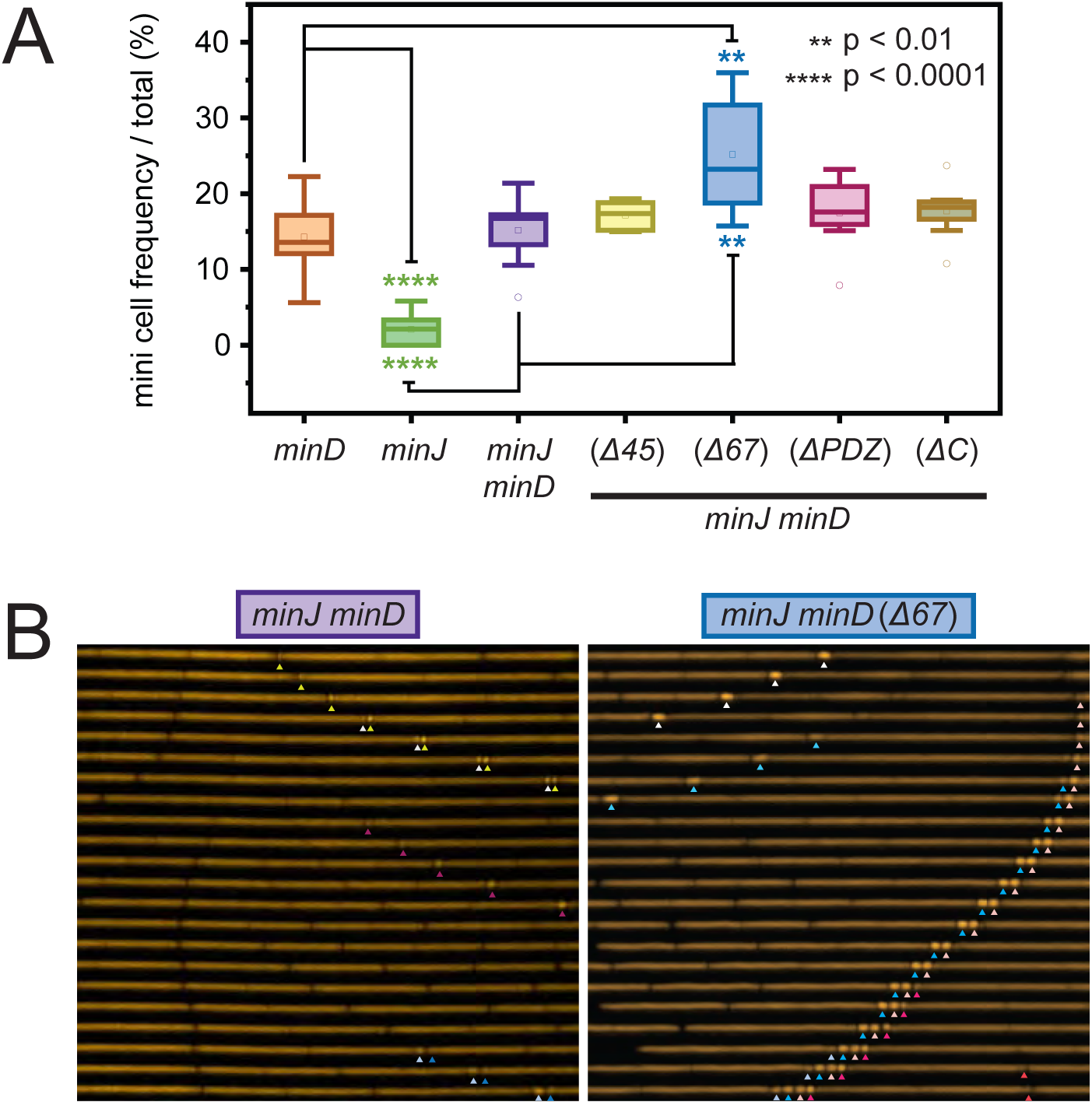
Deletion of the MinJ putative β-sheet results in a hyperminicell phenotype. **A)** Quantification of minicell frequency from cells grown in microfluidic channels expressing a cytoplasmic *P_hyspank_*-mCherry reporter from an ectopic chromosomal locus. Minicell frequency was calculated by dividing the number of minicell-producing division events by the total number of division events observed and multiplying by 100. Each point represents the minicell frequency measured from an individual microfluidic channel. Boxes indicate the distribution of ten independent channels analyzed per strain, the central line indicates the mean, and error bars indicate standard deviation. **B)** Representative kymographs of a microfluidic channel containing a minJ minD and *minD minJ (P_minJ_-minJ^Δ67^)* both expressing *P_hyspank_*-mCherry (false colored yellow). Time progresses from top to bottom at 16-minute intervals over a total period of 336 minutes. Carets indicate minicells generated during successive rounds of cell division. The following strains were used to generate these data: *minD* (DB3728), *minJ* (DB3754), *minD minJ* (DB3639), *minD minJ (P_minJ_-minJ^Δ45^)* (DB3641), *minD minJ (P_minJ_-minJ^Δ67^)* (DB3643), *minD minJ (P_minJ_-minJ^ΔPDZ^)* (DB3642), and *minD minJ (P_minJ_-minJ^ΔC^)* (DB3755).

## DISCUSSION

MinJ is a structurally complex membrane protein and *in silico* topological programs gave divergent predictions as to the number of transmembrane segments. Here, we empirically test for MinJ topology with a classical fusion-based assay and find that the results are most consistent with a 9-pass transmembrane protein that places the N-terminus in the extracellular space and the C-terminal PDZ domain in the cytoplasm. While the PDZ domain was required for MinJ function, it was dispensible for polar localization and may constitute a regulatory motif. Serial deletion of transmembrane segments that preserved overall MinJ topology indicated that each pair was also required MinJ activity, with the deletion of segments 6 and 7 representing the best candidate for interpretation as it produced protein at levels comparable to the wild type. Subsequent analysis indicated that segments 6 and 7 form a putative transmembrane β-sheet while required for subcellular localization and MinJ interaction with itself, MinD, and DivIVA, conferred a change-of-function hyper-minicell phenotype when deleted. We conclude that the bulk of MinJ activity is mediated by the transmembrane domain, and that the function of MinJ is as complex as its structure,

The structure of MinJ is unusual in several regards. For example, the initial transmembrane alpha helix is oriented such that the N-terminus is extracellular, suggesting that membrane insertion does not proceed according to canonical Sec secretion (32–33). More significantly, transmembrane prediction programs failed to correctly identify the nine transmembrane segments, likely due to the fact that two of the segments may form a β-sheet. The β-sheet was predicted by Alphafold3 and is supported by enzyme-fusion based topological analysis, where either side of the sheet is definitively internal, and fusions in the extracellular loop act as though it is both internal and external, an anomaly perhaps consistent with the inability to fully form the sheet. β-sheet transmembrane segments are common in the outer membrane of Gram-negative bacteria where they form the structure of open channel porins (34–35) but are rare to the point of obscurity in the plasma membrane. Indeed, if the β-sheet can be structurally confirmed, MinJ may be the first example of such a protein in the plasma membrane.

The putative transmembrane β-sheet is consistently predicted in all conserved MinJ homologs, and deletion of the sheet (MinJ^Δ67^) was the only mutation in the transmembrane domain that was produced at wild type levels. The MinJ^Δ67^ deletion resulted in a defect in midcell septation that gave rise to long filaments and phenocopied a *minJ* null mutation. Filamentation in the absence of MinJ is thought to be due to the inability to restrict MinCD-mediated depolymerization of FtsZ protofilaments (20–21). Seemingly consistent with an inability to restrict MinD activity, the MinJ^Δ67^ mutant also failed to localize to the cell poles and failed to interact with itself and both DivIVA and MinD, again suggesting a loss-of-function phenotype. Unlike a MinJ null mutation however, deletion of segments 6 and 7 in the absence of MinD resulted in a hyper-minicell phenotype where the frequency of minicell formation exceeded that observed in the *minD* mutant alone. We note that the hyper-minicell phenotype may have been observed in a prior deletion analysis of MinJ, and in that paper it was proposed that MinJ also disassembles the divisome components for peptidoglycan synthesis after septation (23). Perhaps, MinJ and its putative transmembrane β-sheet somehow links the disassembly of the cytoplasmic FtsZ-ring and the membrane-bound divisome.

The MinJ cytoplasmic PDZ domain is also required for activity but how it is required is unknown. For example, deletion of the PDZ domain gives rise to filamentous cells that resemble a *minJ* null mutant, but the resulting protein still exhibits polar localization. Thus, localization of the transmembrane domain alone is insufficient for MinJ activity. Consistent with localization, the MinJ^ΔPDZ^ allele was proficient for interaction with the polar determinant DivIVA when co-expressed in an *E. coli*-based bacterial two-hybrid assay, and was also proficient for interaction with MinD, in the same system. Deletion of the entire C-terminus that includes the PDZ domain showed similar interaction patterns as deletion of the PDZ deletion alone but with seemingly stronger interactions, suggesting perhaps that the C-terminus and PDZ domain could possibly be inhibitory on MinJ activity. As MinJ and MinD localize at the nascent division plane, the C-terminus may represent a regulatory domain to prevent divisome disassembly prior to septum completion.

MinJ is restricted to the rod-shaped Bacillota family of Gram-positive bacteria, and our topological and genetic analysis helps to understand the complex role of MinJ in cell division control. Rather than being a structural nexus, the PDZ domain seems to serve as a regulatory input on the activity of a complex nucleated by the transmembrane domain, and interactors other than MinD and DivIVA have been proposed (24, 27). Moreover, the transmembrane domain is elaborate, perhaps to coordinate timing of disassembly of the internal and external components of the divisome and promote the recycling of each for the next round of cell division. Whatever the case, the MinJ structure seems to be very highly conserved, including all nine transmembrane segments, and is co-conserved with the proteins MinD and DivIVA with which it interacts. The most enigmatic structural element must be the putative transmembrane β-sheet, a motif normally excluded from the cytoplasmic membrane, and mutation of which confers a hyper-minicell phenotype. If the β-sheet can be structurally confirmed, it seems likely to represent a specialized interaction point for cell division regulation, with either another transmembrane protein, or perhaps, a lipid.

## MATERIALS AND METHODS

### Strains and growth conditions

*B. subtilis or E. coli* strains were grown in Luria-Bertani (LB) (10 g tryptone, 5 g yeast extract, 5 g NaCl per L) broth or on LB plates fortified with 1.5% Bacto agar at 37°C. When appropriate, antibiotics were included at the following concentrations: 10 µg/ml tetracycline, 100 µg/ml spectinomycin, 5 µg/ml chloramphenicol, 5 µg/ml kanamycin (*B. subtilis*) or 25 µg/ml kanamycin (*E. coli*), and 1 µg/ml erythromycin plus 25 µg/ml lincomycin (*mls*). Isopropyl b-D-thiogalactopyranoside (IPTG, Sigma) was added to the medium at the indicated concentration when appropriate.

### Swarm expansion assay

Cells were grown overnight at room temperature in LB broth, backdiluted and grown to mid-log phase at 37°C in LB broth and 1mM IPTG (if applicable), and resuspended to an OD_600_ of 10 in MQ H_2_O containing 0.5% India ink. Freshly prepared LB plates containing 0.7% Bacto agar (25 ml/plate) and 1mM IPTG (if applicable) were dried for 10 minutes in a laminar flow hood, centrally inoculated with 10 µl of the cell suspension, dried for another 10 minutes, and incubated at 37°C for 6 hours. Each strain was done in technical triplicate. The India ink demarcates the origin of the inoculation, and the swarm radius was measured in mm relative to this origin. For consistency, an axis was drawn on the back of the plate and swarm radii measurements were taken along this axis.

### β-Galactosidase assays

Three independent biological replicates of each *B. subtilis* strain were grown in 3 mL of LB broth at 37°C to an OD_600_ of 0.7–1.0. Two hundred microliters of cells were harvested and resuspended in 200 µL of Z-buffer (16.1 g Na_2_HPO_4_•7H_2_O + 5.5 g NaH_2_PO_4_•H_2_O + 0.75 g KCl + 0.246 g MgSO_4_•7H_2_O in 1 L H_2_O, pH 7) with 0.27% βME (β-mercaptoethanol). Four microliters of 10 mg/mL lysozyme were added, and cells were lysed by incubating samples at 30°C for 15 min. For the assay, 40 µL of the resuspended cells were transferred to wells of a Greiner-Costar 96-well microtiter plate containing 160 µL Z-buffer, for a final volume of 200 µL per well. Three wells containing 200 µL Z-buffer alone were included as negative controls. Forty microliters of 4 mg/mL ortho-nitrophenyl-β-D-galactopyranoside (ONPG) in Z-buffer were added to each reaction. The plate was incubated at 30°C for 1 h, and the OD_420_ and OD_550_ of each well were taken every 2 min. The slope of each sample’s ODs over time was derived. The average slope of the three negative control wells was subtracted from the slope of each experimental well. The Miller Units were calculated using the following formula. Miller Units = [((slope of OD₄₂₀) − 1.75 × (slope of OD₅₅₀)) / (lysate volume (mL) × OD₆₀₀)] × 1000 × dilution factor

### Alkaline phosphatase assays

Three independent biological replicates of each *B. subtilis* strain were grown in 3 mL of LB broth at 37°C to an OD_600_ of 0.7–1.0. Two hundred microliters of cells were harvested by centrifugation and resuspended in 200 µL Tris-buffer (pH 8). For the assay, 40 µL of the resuspended cells were transferred to wells of a Greiner-Costar 96-well microtiter plate containing 160 µL Tris-buffer, for a final volume of 200 µL per well. Three wells containing 200 µL Tris-buffer alone were included as negative controls. Forty microliters of 4 mg/mL p-Nitrophenyl Phosphate (pNPP) in Tris-buffer was added to each reaction. The plate was incubated at 37°C for 2 h, and the OD_420_ and OD_550_ of each well were taken every 2 min. The slope of each sample’s ODs over time was derived. The average slope of the three negative control wells was subtracted from the slope of each experimental well. The Miller Units were calculated using the following formula. Miller Units = [((slope of OD₄₂₀) − 1.75 × (slope of OD₅₅₀)) / (lysate volume (mL) × OD₆₀₀)] × 1000 × dilution factor

### Microscopy

Strains of *B. subtilis* were grown in LB (with antibiotics or IPTG as indicated) at 37℃ to an exponential OD_600_ (0.4-0.8). For membrane visualization, cells were harvested by centrifugation and resuspended in 30 µL of MQH_2_O containing 15 µg/ml of FM4-64 (Invitrogen #T13320). Samples were incubated for 2 min at room temperature in the dark and washed twice before imaging. For imaging, 4 µL of the stained cell suspension was spotted onto a glass microscope slide and immobilized using a poly-L-lysine-coated coverslip. Fluorescence microscopy was performed using a Nikon Ti2E inverted microscope equipped with a Plan Apo 100×/1.4 NA oil immersion phase-contrast objective and a Prime 95B camera. Images were captured using NIS-Elements software and processed using Fiji/ImageJ software.

### Western Blotting

*B. subtilis* strains were grown in LB to an OD_600_ range of 0.7-0.8, 1 ml was harvested by centrifugation, resuspended to 10 OD_600_ in Lysis buffer (20 mM Tris pH 7.0, 1 mg/ml lysozyme, 10 mg/ml DNAse I, 100 mg/ml RNAse I, 1 mM PMSF, 10 mg/ml MgCl_2_) and incubated 30 min at 37°C. 10 ml of lysate was mixed with 2 ml 6x SDS loading dye. Samples were separated by 12% Sodium dodecyl sulfate-polyacrylamide gel electrophoresis (SDS-PAGE). The proteins were electroblotted onto nitrocellulose and developed with a 1:80,000 dilution of anti-SigA or 1:5,000 dilution of anti-MinJ^PDZ^, and a 1:10,000 dilution secondary antibody (horseradish peroxidase-conjugated goat anti-rabbit immunoglobulin G). Immunoblot was developed using the Immun-Star HRP developer kit (Bio-Rad).

### Bacterial-Two-Hybrid Beta-galactosidase assay

This assay is based on the single step method for β-galactosidase (36). Three independent biological replicates of each *E. coli* strain were grown in 3 mL of LB broth at 37°C to an OD_600_ of ∼ 1.0. Subsequently, 20ul of each culture was inoculated into 3 mL of fresh LB broth and grown overnight at room temperature (25°C) to an OD_600_ of ∼1.0. For β-galactosidase measurements, 120 µL of reaction mixture (Z-buffer supplemented with 0.27% β-mercaptoethanol (βME), 4 mg/mL ONPG, 80 µL PopCulture reagent, and 10 mg/mL lysozyme) was dispensed into wells of a Greiner-Costar 96-well plate. Eighty microliters of cells were added directly to the reaction mixture in each well, and 200 µL of reaction mixture alone were included in triplicate as negative controls. The plate was incubated in a plate reader for 1 h at 28°C, with double orbital continuous shake, and the OD_420_ and OD_600_ of each well were taken every 1 min. The slope of each sample’s OD_420_ over time was derived. The average slope of the three negative control wells was subtracted from the slope of each experimental well. The Miller Units were calculated using the following formula: Miller Units = 5000 × (OD₄₂₀ slope) / Initial OD₆₀₀

### SPP1 phage transduction

To 0.2 mL of dense culture grown in TY broth (LB broth supplemented after autoclaving with 10 mM MgSO_4_ and 100 µM MnSO_4_), serial dilutions of SPP1 phage stock were added and statically incubated for 15 min at 37°C. To each mixture, 3 mL TYSA (molten TY supplemented with 0.5% agar) was added, poured atop fresh TY plates, and incubated at 30°C overnight. Top agar from the plate containing near confluent plaques was harvested by scraping into a 50 ml conical tube, vortexed, and centrifuged at 5,000 x g for 10 min. The supernatant was treated with 25 µg/mL DNase final concentration before being passed through a 0.45 µm syringe filter and stored at 4°C.

Recipient cells were grown to stationary phase in 3 mL TY broth at 37°C. 1 ml cells were mixed with 25 µL of SPP1 donor phage stock. Nine mL of TY broth was added to the mixture and allowed to stand at 37°C for 20 min. The transduction mixture was then centrifuged at 5,000 x g for 10 min, the supernatant was discarded and the pellet was resuspended in the remaining volume. Cell suspension (100 µL) was then plated on TY fortified with 1.5% agar, the appropriate antibiotic, and 10 mM sodium citrate.

### Microfluidics

#### Materials

Materials used to fabricate microfluidics devices included: glass slides (25 mm by 75 mm) and No. 1.5 cover glass (24 mm x 60 mm) from VWR International, LLC; poly(dimethylsiloxane) (PDMS, Sylgard 184) from Dow Corning, Inc.; SU-8 2010 photoresist and SU-8 developer from Kayaku Advanced Materials, Inc.; titanium di-isopropoxide bis(2,4-pentanedionate) from Gelest, Inc.; and all other chemicals from Sigma-Aldrich Co.

#### Device Fabrication and Operation

A new simplified microfluidic device developed by Joncha (Unpublished) consisted of three layers: cover glass (bottom), a ∼100-µm thick PDMS fluid layer (middle), and a 3-mm thick PDMS control layer (top). The PDMS layers were fabricated through a mold-replica process (14, 37). The SU-8 mold for the fluid layer was formed in two steps: electron beam lithography (FEI Quanta 600F scanning electron microscope equipped with a JC Nabity Nanometer Pattern Generation System) generated arrays of approximately 1000 microchannels (1.0 µm deep and 1.0 µm wide) followed by photolithography (OAI 200 Mask Aligner) to pattern microchannels (20 µm deep, 60 µm wide, and 200 µm center-to-center) positioned perpendicularly to the microchannel arrays. A 10:1 ratio of PDMS polymer to cross-linker was spin-coated onto the fluid-layer mold at 1500 rpm to achieve an ∼100-µm thick layer and poured onto the control-layer mold before being degassed. The PDMS for the fluid layer was partially cured, whereas the PDMS for the control layer was fully cured. The control layer was removed from its mold and aligned onto the fluid layer, which was then cured in a 75°C oven for at least 4 h to fully bond the two layers. The side of the fluid layer with the microchannel arrays was plasma cleaned (Harrick PDC-001) and bonded to the cover glass. Microchannels were filled with LB media with 0.1% bovine serum albumin, and the device was checked for any defects. Media was fed through the device by gravity. Cells were loaded and trapped in the microchannel arrays by applying pressure from above. Cells were allowed to grow and acclimate for approximately 2 h before analysis began.

#### Time-Lapse Image Acquisition

Fluorescence images were collected on a Nikon Ti2 microscope equipped with a Hamamatsu ORCA-Fusion Digital CMOS camera, 100X Plan Apo oil-immersion objective (1.45 NA), and Lumencor SPECTRA light engine. The mCherry fluorophore was excited at 554/23 nm, and emission was collected at 595/31 nm with an exposure time of 250 ms. The objective warmer (Okolab) was set at 30°C for all experiments. Nikon NIS-Elements was used for brightfield and mCherry image acquisition at 2-min intervals at 10 locations across the microchannel arrays. Denoise.ai mode was used to decrease noise due to stage jitter.

#### Data Analysis

Image files in ND2 format were parsed into 10 separate files based on the 10 locations imaged across the microchannel arrays. With Fiji software (38), horizontal positions of the images were adjusted to 0° to track cell position over time, and backgrounds were subtracted from each set of images. If images shifted laterally over time, time-lapse sequences were realigned with the SIFT multichannel function (39) in the Fiji software. Image sequences of individual channels containing viable cells were generated from images taken at 2-min intervals. To identify both cells and minicells, localization of mCherry (cytoplasm) was automatically tracked with the MicrobeJ tracking plug-in (40), and cell identification was manually confirmed. To differentiate cells from minicells, length thresholds were set at 4 to 50 µm for elongated/dividing cells and 0.1 to 2 µm for mini cells. After automatic tracking and manual filtering, mini cell frequency was calculated from the count of minicells over the total number of cells (elongated/division cells plus minicells).

### Bioinformatic Analysis

#### Phylogenetic Reconstruction and Co-occurrence Analysis

To assess the phylogenetic distribution of MinJ across the tree of life, representative species spanning Bacteria, Archaea, and Eukarya were selected, with bacterial representatives broadly sampling the major phyla. NCBI taxonomic identifiers (TaxIDs) for each species were compiled and used to query the NCBI Protein database for the presence of MinJ. Species were scored as MinJ-positive when at least one annotated MinJ protein was detected. Domain and phylum information for each species were retrieved from the NCBI taxonomy database and used to generate a tree of life using PhyloT (https://phylot.biobyte.de/). The resulting tree was visualized and annotated using iTOL(41). Following the observation that MinJ was restricted to members of the Firmicutes, the MinJ protein sequence from *Bacillus subtilis* NCIB3610 was used to identify orthologs through EggNOG (42), yielding 396 MinJ orthologs. Protein sequences were retrieved in FASTA format, aligned using MAFFT, and trimmed prior to phylogenetic reconstruction. Maximum-likelihood phylogenetic inference was performed using IQ-TREE with ModelFinder enabled for automatic model selection. The best-fitting amino acid substitution model was LG+F+R8. The resulting phylogenetic tree was visualized and annotated using iTOL.

To investigate the evolutionary association of MinJ with the cell division proteins MinD and DivIVA, species represented within the MinJ ortholog dataset were queried using their NCBI TaxIDs. Presence or absence of MinD and DivIVA homologs was determined through NCBI Protein database searches and recorded as binary values, where 1 indicated presence and 0 indicated absence. These data were compiled into binary matrices and imported into iTOL as annotation datasets to visualize the distribution of MinD and DivIVA across the MinJ phylogeny.

#### Homology and Structural Analysis

To identify conserved residues within MinJ, the multiple sequence alignment generated from the MinJ ortholog dataset was analyzed using custom Python scripts implemented with Biopython. For each alignment position, the most frequently occurring amino acid residue was identified and the percentage conservation calculated across all aligned sequences. Residues were subsequently ranked according to their degree of conservation and Shannon entropy. Thereafter, highly conserved residues were identified by selecting positions exhibiting greater than 90% conservation and mapping them onto the *Bacillus subtilis* MinJ sequence. To identify conserved motifs, contiguous stretches of residues exhibiting at least 50% conservation across orthologs were identified and mapped onto the reference sequence. The average conservation of each conserved stretch was then calculated.

To assess structural conservation of MinJ, the AlphaFold-predicted structure of *Bacillus subtilis* MinJ (UniProt accession O34775) was used as a query for structural similarity searches using Foldseek (43). Subsequently, structural homologs identified by Foldseek were examined for their phylogenetic distribution and used to assess conservation of the MinJ fold across bacterial lineages. To further investigate structural divergence within the MinJ family, representative MinJ orthologs were selected from progressively more distant branches of the MinJ phylogeny and the amino acid sequences from these representatives were submitted to AlphaFold3 for structure prediction. Predicted structures were subsequently compared to assess conservation of overall architecture, transmembrane organization, and structural features identified in the *Bacillus subtilis* MinJ model.

### Strains Construction

All constructs were either first introduced into the domesticated strain PY79 by natural competence and then transferred to the 3610 background using SPP1-mediated generalized phage transduction (44), or by direct transformation into the competent derivative of 3610, DK1042 (45). All strains used in this study are listed in **Table 1**. All plasmids used in this study are listed in **Table S3**. All primers used in this study are listed in **Table S4**.

**Table 1.** Bacillus subtilis strains.

| Strains | Genotype |
| --- | --- |
| DB92 | *ComI*<sup>Q12L</sup> *minJ*::*tet* |
| DB2484 | *ComI*<sup>Q12L</sup> *amyE*::*P*<sub>*minJ*</sub>-*minJ spec minJ*::*tet* |
| DB2485 | *ComI*<sup>Q12L</sup> *amyE*::*P*<sub>*minJ*</sub>-*minJ*<sup>ΔC</sup> *spec minJ*::*tet* |
| DB2499 | *ComI*<sup>Q12L</sup> *amyE*::*P*<sub>*minJ*</sub>-*minJ*<sup>Δ23</sup> *spec minJ*::*tet* |
| DB2500 | *ComI*<sup>Q12L</sup> *amyE*::*P*<sub>*minJ*</sub>-*minJ*<sup>Δ45</sup> *spec minJ*::*tet* |
| DB2501 | *ComI*<sup>Q12L</sup> *amyE*::*P*<sub>*minJ*</sub>-*minJ*<sup>Δ89</sup> *spec minJ*::*tet* |
| DB2502 | *ComI*<sup>Q12L</sup> *amyE*::*P*<sub>*minJ*</sub>-*minJ*<sup>ΔPDZ</sup> *spec minJ*::*tet* |
| DB2968 | *ComI*<sup>Q12L</sup> *amyE*::*P*<sub>*minJ*</sub>-*minJ-mneongreen spec minJ*::*tet* |
| DB3144 | *ComI*<sup>Q12L</sup> *amyE*::*P*<sub>*minJ*</sub>-*minJ*<sup>Δ67</sup> *spec minJ*::*tet* |
| DB3506 | *ComI*<sup>Q12L</sup> *amyE*::*P*<sub>*minJ*</sub>-*minJ-mneongreen spec minD*::*cat minJ*::*tet* |
| DB3508 | *ComI*<sup>Q12L</sup> *amyE*::*P*<sub>*minJ*</sub>-*minJ*<sup>Δ67</sup>-*mneongreen spec minD*::*cat minJ*::*tet* |
| DB3509 | *ComI*<sup>Q12L</sup> *amyE*::*P*<sub>*minJ*</sub>-*minJ*<sup>ΔPDZ</sup>-*mneongreen spec minD*::*cat minJ*::*tet* |
| DB3582 | *ComI*<sup>Q12L</sup> *amyE*::*P*<sub>*minJ*</sub>-*minJ*<sup>Δ45</sup>-*mneongreen spec minD*::*cat minJ*::*tet* |
| DB3626 | *ComI*<sup>Q12L</sup> *amyE*::*P*<sub>*minJ*</sub>-*minJ*<sup>ΔC</sup>-*mneongreen spec minD*::*cat minJ*::*tet* |
| DB3639 | *ComI*<sup>Q12L</sup> *minJ*::*tet minD*::*TnYLB kan thrC*::*P*<sub>*hyspank*</sub>-*mcherry mls* |
| DB3641 | *ComI*<sup>Q12L</sup> *amyE*::*P*<sub>*minJ*</sub>-*minJ*<sup>Δ45</sup> *spec minD*::*cat minJ*::*tet thrC*::*P*<sub>*hyspank*</sub>-*mcherry mls* |
| DB3642 | *ComI*<sup>Q12L</sup> *amyE*::*P*<sub>*minJ*</sub>-*minJ*<sup>ΔPDZ</sup> *spec minD*::*cat minJ*::*tet thrC*::*P*<sub>*hyspank*</sub>-*mcherry mls* |
| DB3643 | *ComI*<sup>Q12L</sup> *amyE*::*P*<sub>*minJ*</sub>-*minJ*<sup>Δ67</sup> *spec minD*::*cat minJ*::*tet thrC*::*P*<sub>*hyspank*</sub>-*mcherry mls* |
| DB3728 | *ComI*<sup>Q12L</sup> *amyE*::*P*<sub>*minJ*</sub>-*minJ spec minD*::*cat minJ*::*tet thrC*::*P*<sub>*hyspank*</sub>-*mcherry mls* |
| DB3754 | *ComI*<sup>Q12L</sup> *minJ*::*tet thrC*::*P*<sub>*hyspank*</sub>-*mcherry mls* |
| DB3755 | *ComI*<sup>Q12L</sup> *amyE*::*P*<sub>*minJ*</sub>-*minJ*<sup>ΔC</sup> *spec minD*::*cat minJ*::*tet thrC*::*P*<sub>*hyspank*</sub>-*mcherry mls* |
| DK1042 | *ComI*<sup>Q12L</sup> |
| DK9494 | *amyE*::*P*<sub>*hyspank*</sub>-*minJ*<sup>@240</sup>-*lacZ spec* |
| DK9495 | *amyE*::*P*<sub>*hyspank*</sub>-*minJ*<sup>@305</sup>-*lacZ spec* |
| DK9580 | *amyE*::*P*<sub>*hyspank*</sub>-*minJ*<sup>@167</sup>-*lacZ spec* |
| DK9581 | *amyE*::*P*<sub>*hyspank*</sub>-*minJ*<sup>@132</sup>-*lacZ spec* |
| DK9582 | *amyE*::*P*<sub>*hyspank*</sub>-*minJ*<sup>@99</sup>-*lacZ spec* |
| DK9583 | *amyE*::*P*<sub>*hyspank*</sub>-*minJ*<sup>@49</sup>-*lacZ spec* |
| DK9623 | *phoB*::*tet phoA*::*mls amyE*::*P*<sub>*hyspank*</sub>-*minJ*<sup>@305</sup>-*phoA spec* |
| DK9624 | *phoB*::*tet phoA*::*mls amyE*::*P*<sub>*hyspank*</sub>-*minJ*<sup>@240</sup>-*phoA spec* |
| DK9625 | *phoB*::*tet phoA*::*mls amyE*::*P*<sub>*hyspank*</sub>-*minJ*<sup>@167</sup>-*phoA spec* |
| DK9626 | *phoB::tet phoA::mls amyE::P<sub>hyspank</sub>-minJ@132-phoA spec* |
| DK9627 | *phoB::tet phoA::mls amyE::P<sub>hyspank</sub>-minJ@99-phoA spec* |
| DK9628 | *phoB::tet phoA::mls amyE::P<sub>hyspank</sub>-minJ@49-phoA spec* |
| DK9668 | *amyE::P<sub>hyspank</sub>-minJ@209-lacZ spec* |
| DK9669 | *amyE::P<sub>hyspank</sub>-minJ@268-lacZ spec* |
| DK9670 | *phoB::tet phoA::mls amyE::P<sub>hyspank</sub>-minJ@209-phoA spec* |
| DK9671 | *phoB::tet phoA::mls amyE::P<sub>hyspank</sub>-minJ@268-phoA spec* |
| DK9791 | *amyE::P<sub>hyspank</sub>-minJ@395-lacZ spec* |
| DK9804 | *amyE::P<sub>hyspank</sub>-minJ@197-lacZ spec* |
| DK9866 | *amyE::P<sub>hyspank</sub>-minJ@81-lacZ spec* |
| DK9872 | *phoB::tet phoA::mls amyE::P<sub>hyspank</sub>-minJ@81-phoA spec* |
| DK9873 | *phoB::tet phoA::mls amyE::P<sub>hyspank</sub>-minJ@197-phoA spec* |
| DK9874 | *phoB::tet phoA::mls amyE::P<sub>hyspank</sub>-minJ@395-phoA spec* |

**Table 2.** *E.coli* strains.

| Strains | Genotype |
| --- | --- |
| DE4545 | *minJ-linker-cyaAT18 amp, minJ-linker-cyaAT25 kan* |
| DE4546 | *minJ<sup>Δ45</sup> -linker-cyaAT18 amp, minJ<sup>Δ45</sup> -linker-cyaAT25 kan* |
| DE4547 | *minJ<sup>Δ67</sup> -linker-cyaAT18 amp, minJ<sup>Δ67</sup> -linker-cyaAT25 kan* |
| DE4548 | *minJ<sup>ΔPDZ</sup> -linker-cyaAT18 amp, minJ<sup>ΔPDZ</sup> -linker-cyaAT25 kan* |
| DE4550 | *minJ<sup>ΔC</sup> -linker-cyaAT18 amp, minJ<sup>ΔC</sup> -linker-cyaAT25 kan* |
| DE4552 | *minJ-linker-cyaAT18 amp, minD-linker-cyaAT25 kan* |
| DE4553 | *minJ<sup>Δ45</sup> -linker-cyaAT18 amp, minD-linker-cyaAT25 kan* |
| DE4554 | *minJ<sup>Δ67</sup> -linker-cyaAT18 amp, minD-linker-cyaAT25 kan* |
| DE4555 | *minJ<sup>ΔPDZ</sup> -linker-cyaAT18 amp, minD-linker-cyaAT25 kan* |
| DE4557 | *minJ<sup>ΔC</sup> -linker-cyaAT18 amp, minD-linker-cyaAT25 kan* |
| DE4559 | *minJ-linker-cyaAT18 amp, divIVA-linker-cyaAT25 kan* |
| DE4560 | *minJ<sup>Δ45</sup> -linker-cyaAT18 amp, divIVA-linker-cyaAT25 kan* |
| DE4561 | *minJ<sup>Δ67</sup> -linker-cyaAT18 amp, divIVA-linker-cyaAT25 kan* |
| DE4562 | *minJ<sup>ΔPDZ</sup> -linker-cyaAT18 amp, divIVA-linker-cyaAT25 kan* |
| DE4564 | *minJ<sup>ΔC</sup> -linker-cyaAT18 amp, divIVA-linker-cyaAT25 kan* |
| DE4566 | *cyaAT18 amp, cyaAT25 kan* |

#### Topology system constructs

To generate the inducible *minJ*-phoA/lacZ translational reporter fusions, a series of progressively truncated *minJ* fragments was designed based on bioinformatic predictions of membrane topology from TMPRED (46), DEEPTMHMM (47), Uniprot (48) and AlphaFold3 (49). Truncations terminating at amino acid residues aa395, aa305, aa268, aa240, aa209, aa197, aa167, aa132, aa99, aa81 and aa49 were PCR-amplified from *B. subtilis* 3610 chromosomal DNA using primer pairs 7642/7731, 7642/7644, 7642/7713, 7642/7643, 7642/7712, 7642/7730, 7642/7675, 7642/7676, 7642/7677, 7642/7729, and 7642/7678, respectively. The resulting PCR products were digested with SalI/NheI and cloned into the same restriction sites of pDP545 (30), containing the *P_hyspank_*promoter upstream of the *phoA* reporter gene, generating in-frame C-terminal translational fusions in which each *minJ* truncation was positioned downstream of the inducible promoter and immediately upstream of the phoA sequence. The resulting constructs corresponded to plasmids pKO30 (*minJ^@395^*-PhoA), pKO5 (*minJ^@305^*-PhoA), pKO21 (*minJ^@268^*-PhoA), pKO3 (*minJ^@240^*-PhoA), pKO20 (*minJ^@209^*-PhoA), pKO28 (*minJ^@197^*-PhoA), pKO12 (*minJ^@167^*-PhoA), pKO13 (*minJ^@132^*-PhoA), pKO14 (*minJ^@99^*-PhoA), pKO26 (*minJ^@81^*-PhoA), and pKO15 (*minJ^@49^*-PhoA)

The same PCR products were digested with SalI/NheI and cloned into the same restriction sites of pDP559 (Phillips 2021), containing the *P_hyspank_* promoter upstream of the *lacZ* reporter gene, generating in-frame C-terminal translational fusions in which each *minJ* truncation was positioned downstream of the inducible promoter and immediately upstream of the LacZ sequence. The resulting constructs corresponded to plasmids pKO29 (*minJ^@395^*-*lacZ*), pKO4 (*minJ^@305^*-*lacZ*), pKO19 (*minJ^@268^*-*lacZ*), pKO2 (*minJ^@240^*-*lacZ*), pKO18 (*minJ^@209^*-*lacZ*), pKO27 (*minJ*^@197^-*lacZ*), pKO8 (*minJ^@167^*-*lacZ*), pKO9 (*minJ^@132^*-*LacZ*), pKO10 (*minJ^@99^*-*lacZ*), pKO25 (*minJ^@81^*-*lacZ*), and pKO11 (*minJ^@49^*-*lacZ*)

#### *P_hyspank_-minJ* inducible constructs

To generate the inducible *minJ* full length construct pKO35, the *minJ* coding sequence was PCR-amplified from *B. subtilis* 3610 chromosomal DNA using primers 7906/7907. The PCR product was digested with SalI/NheI and cloned into the same sites of pDR111 containing a polylinker downstream of the *P_hyspank_* promoter gene, and a spectinomycin resistance cassette between two arms of the *amyE* gene (generous gift of David Rudner, Harvard Medical School).

To generate the inducible *minJ* truncations pKO66 (*minJ^Δ23^*), pKO67 (*minJ^Δ45^*), pKO68 (*minJ^Δ67^*), pKO69 (*minJ^Δ89^*), and pKO70 (*minJ^ΔPDZ^*), the corresponding upstream fragments were amplified from 3610 chromosomal DNA using primer pairs 7906/8471, 7906/8473, 7906/8468, 7906/8477, and 7906/8479, respectively. The corresponding downstream fragments were amplified using primer pairs 8470/7907, 8472/7907, 8469/7907, 8476/7907, and 8478/7907, respectively. For each truncation, the paired upstream and downstream fragments were joined by isothermal assembly (Gibson 2009). The resulting assembled products were then used as templates for PCR amplification with primers 7906/7907 to generate the final truncation inserts. Each PCR product was digested with SalI/NheI and cloned into the same sites of pDR111 containing a polylinker downstream of the *P_hyspank_* promoter gene, and a spectinomycin resistance cassette between two arms of the *amyE* gene.

To generate pKO73 corresponding to the inducible *minJ^ΔC^* construct, the *minJ* gene was PCR amplified from 3610 chromosomal DNA using primer pair 7906/7907. The PCR product was digested with SalI/NheI and cloned into the same sites of pDR111 containing a polylinker downstream of the *P_hyspank_* promoter gene, and a spectinomycin resistance cassette between two arms of the *amyE* gene (generous gift of David Runder, Harvard Medical School).

#### P_minJ_-minJ deletions

To generate the native *minJ* deletion constructs pKO75 (*minJ^Δ23^*), pKO76 (*minJ^Δ45^*), pKO123 (*minJ^Δ67^*), pKO78 (*minJ^Δ89^*), and pKO79 (*minJ^ΔPDZ^*), the corresponding upstream fragments, containing the native *minJ* promoter region, were amplified from *B. subtilis* 3610 chromosomal DNA using primers 8635/8636. The corresponding downstream truncation fragments were amplified from plasmid templates pKO66, pKO67, pKO68, pKO69, and pKO70, respectively, using primers 8637/8638. The integration vector pAH25 (generous gift from Amy Camp, Mount Holyoke College) was linearized with SalI, and each pair of upstream and downstream fragments was assembled into the vector by isothermal assembly (50).

#### P_minJ_-minJ-mNeongreen constructs

An intermediate vector, pKO103, containing an in-frame linker fused to *mNeongreen* (*mNG*), was first constructed by PCR amplification of *mNG* from pDP427 using primers 8844/8845. The resulting PCR product was digested with SalI/EcoRI and cloned into the corresponding sites of pAH25, which contains a polylinker and a spectinomycin-resistance cassette flanked by *amyE* homology arms.

Subsequently, to generate the native-expression P_minJ_-*minJ*-mNeongreen fusion constructs corresponding to full-length pKO110 (*minJ-mNG*) and the truncation derivatives pKO127 (*minJ^Δ45^-mNG*), pKO115 (*minJ^Δ67^-mNG*), and pKO116 (*minJ^ΔPDZ^-mNG*), the full length *minJ* allele or corresponding truncation variants were amplified from plasmid templates pKO72, pKO76, pKO123, and pKO79, respectively, using primers 8846/8847. The vector plasmid pKO103 was linearized with SalI, and each insert was cloned into the vector by isothermal assembly, generating in-frame C-terminal fusions positioned immediately upstream of the linker-mNG sequence.

#### GST-PDZ

To generate the inducible *minJ^PDZ^* cytoplasmic domain for protein purification length construct pKO37, the *minJ^PDZ^*coding sequence was PCR-amplified from *B. subtilis* 3610 chromosomal DNA using primers 7956/7957. The PCR product was digested with BamHI/EcoRI and cloned into the same sites of pGEX-2TK containing GST coding sequence downstream of the *P_tac_* promoter gene, and an ampicillin resistance cassette.

#### P_hyspank_-mCherry

To generate the inducible *mCherry* reporter construct pKO143, the mCherry coding sequence was PCR-amplified off pEV6 using primers 8399/8400. The PCR product was digested with HindIII/SphI and cloned into the same sites of pDP150 containing a *P_hyspank_* promoter gene, and an erythromycin resistance cassette between two arms of the *thrC* gene

#### Bacterial two-hybrid constructs

To generate pKO71 (*minJ*-linker-CyaAT25), the *minJ* coding sequence was amplified from *B. subtilis* 3610 chromosomal DNA using primers 8818/8819. The PCR product was digested with BamHI/EcoRI and cloned into the same sites of pKNT25 containing the T25 fragment of the *Bordetella pertussis* adenylate cyclase (CyaA) downstream of the P_tac_ promoter and a kanamycin-resistance cassette. The resulting construct generated an in-frame fusion in which *minJ* was positioned upstream of the T25 fragment and downstream of the P_tac_ promoter. To generate pKO106 (*minJ*-linker-CyaAT18), the same digested PCR product was cloned into the corresponding sites of pUT18, containing the T18 fragment of *B. pertussis* adenylate cyclase downstream of the P_tac_ promoter and an ampicillin-resistance cassette. This generated an in-frame fusion in which *minJ* was positioned upstream of the T18 fragment and downstream of the P_tac_ promoter.

To generate additional *minJ* deletion alleles for bacterial two-hybrid analysis, primers 8818/8819 were used to amplify the corresponding alleles from plasmid templates pKO76 (*minJ* ^Δ45^), pKO123(*minJ* ^Δ67^), pKO79(*minJ* ^ΔPDZ^). The resulting PCR products were digested with BamHI/EcoR1 and cloned into pKNT25 to generate pKO148 (*minJ* ^Δ45^*-linker-cyaAT25)*, pKO149 (*minJ* ^Δ67^*-linker-cyaAT25)*, pKO136 (*minJ* ^ΔPDZ^*-linker-cyaAT25*) and pKO155 (*minJ* ^ΔC^ *-linker- cyaAT25 kan)*. The same digested products were cloned into pUT18C to generate pKO151 (*minJ* ^Δ45^*-linker-cyaAT18)*, pKO147 (*minJ* ^Δ67^*-linker-cyaAT18)*, pKO146 (*minJ* ^ΔPDZ^*-linker-cyaAT18*) and pKO157 (*minJ* ^ΔC^ *-linker-cyaAT25 amp)*.

## ACKNOWLEDGEMENTS

We thank Cristina Landeta for consultation and advice on *lacZ* and *phoA* based topology analysis, Seyi Esan for help with constructing constructs for the bacterial two-hybrid analysis, and the Nanoscience Core Facility for use of its instruments. The work was funded by NIH R35GM131783 to DBK and NIH R35GM141922 to SCJ.

---

**Figure S1.**
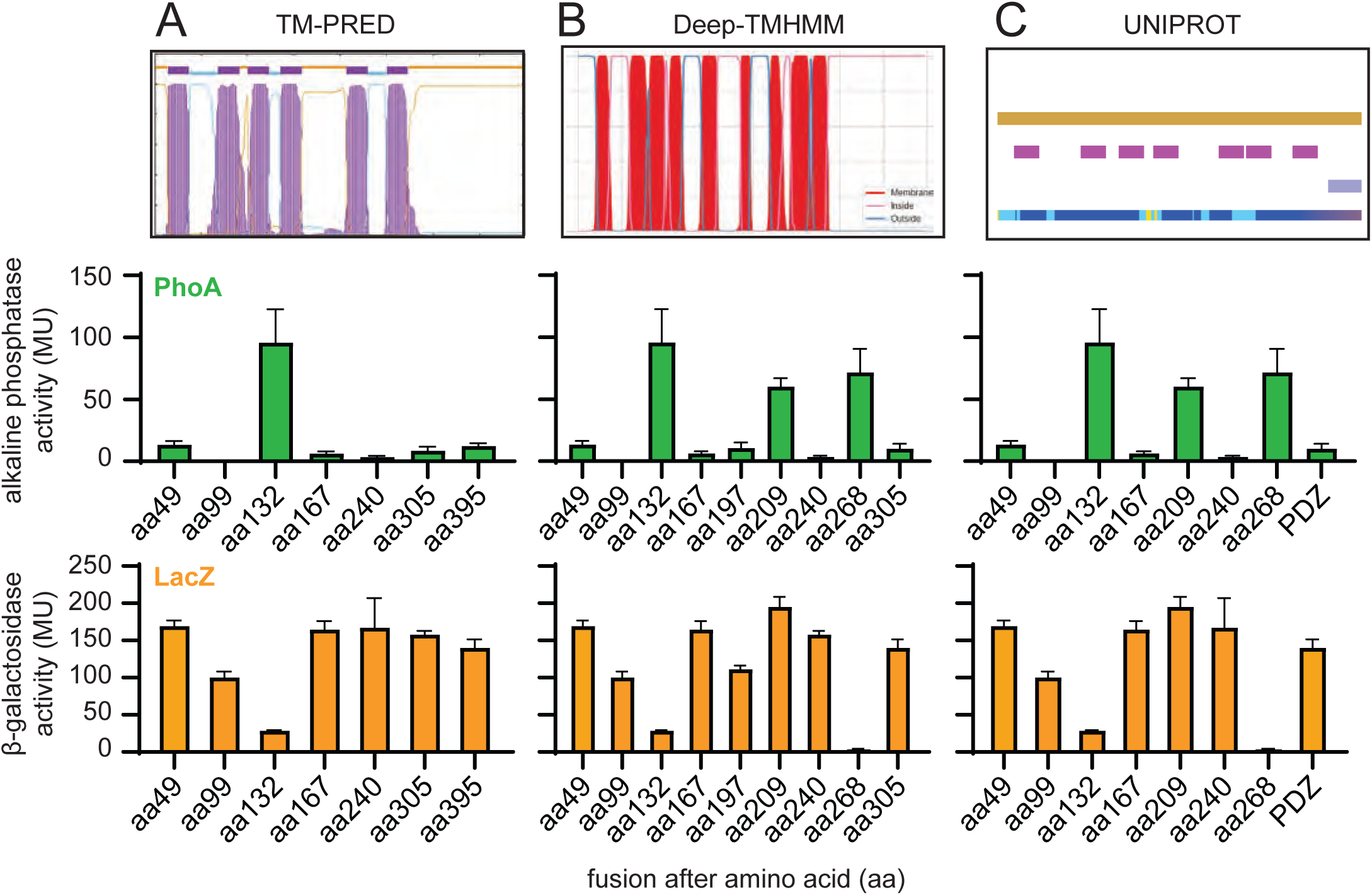
Topological models predicted by TMPRED, DeepTMHMM, and UniProt are inconsistent with experimental topology mapping. A) Topology model generated using TMPRED. The upper panel shows the predicted transmembrane architecture of MinJ. The middle panel shows alkaline phosphatase (PhoA) activity of the corresponding MinJ-PhoA translational fusions, and the lower panel shows β-galactosidase (LacZ) activity of the corresponding MinJ-LacZ translational fusions. B) Topology model generated using DeepTMHMM with corresponding PhoA and LacZ reporter activities shown below. C) Topology model generated using UniProt with corresponding PhoA and LacZ reporter activities shown below. The following strains were used to generate these data: PhoA fusions DK9874 (*minJ^@395^*-PhoA), DK9623 (*minJ^@305^*-PhoA), DK9671 (*minJ^@268^*-PhoA), DK9624 (*minJ^@240^*-PhoA), DK9670 (*minJ^@209^*-PhoA), DK9873 (*minJ^@197^*-PhoA), DK9625 (*minJ^@167^*-PhoA), DK9626 (*minJ^@132^*-PhoA), DK9627 (*minJ^@99^*-PhoA), DK9872 (*minJ^@81^*-PhoA), and DK9628 (*minJ^@49^*-PhoA) and LacZ fusions; DK9791 (*minJ^@395^*- *lacZ*), DK9495 (*minJ^@305^*-*lacZ*), DK9669 (*minJ^@268^*-*lacZ*), DK9494 (*minJ^@240^*-*lacZ*), DK9668 (*minJ^@209^*-*lacZ*), DK9804 (*minJ*^@197^-*lacZ*), DK9580 (*minJ^@167^*-*lacZ*), DK9581 (*minJ^@132^*-*LacZ*), DK9582 (*minJ^@99^*-*lacZ*), DK9866 (*minJ^@81^*-*lacZ*), and DK9583 (*minJ^@49^*-*lacZ*)

**Figure S2.**
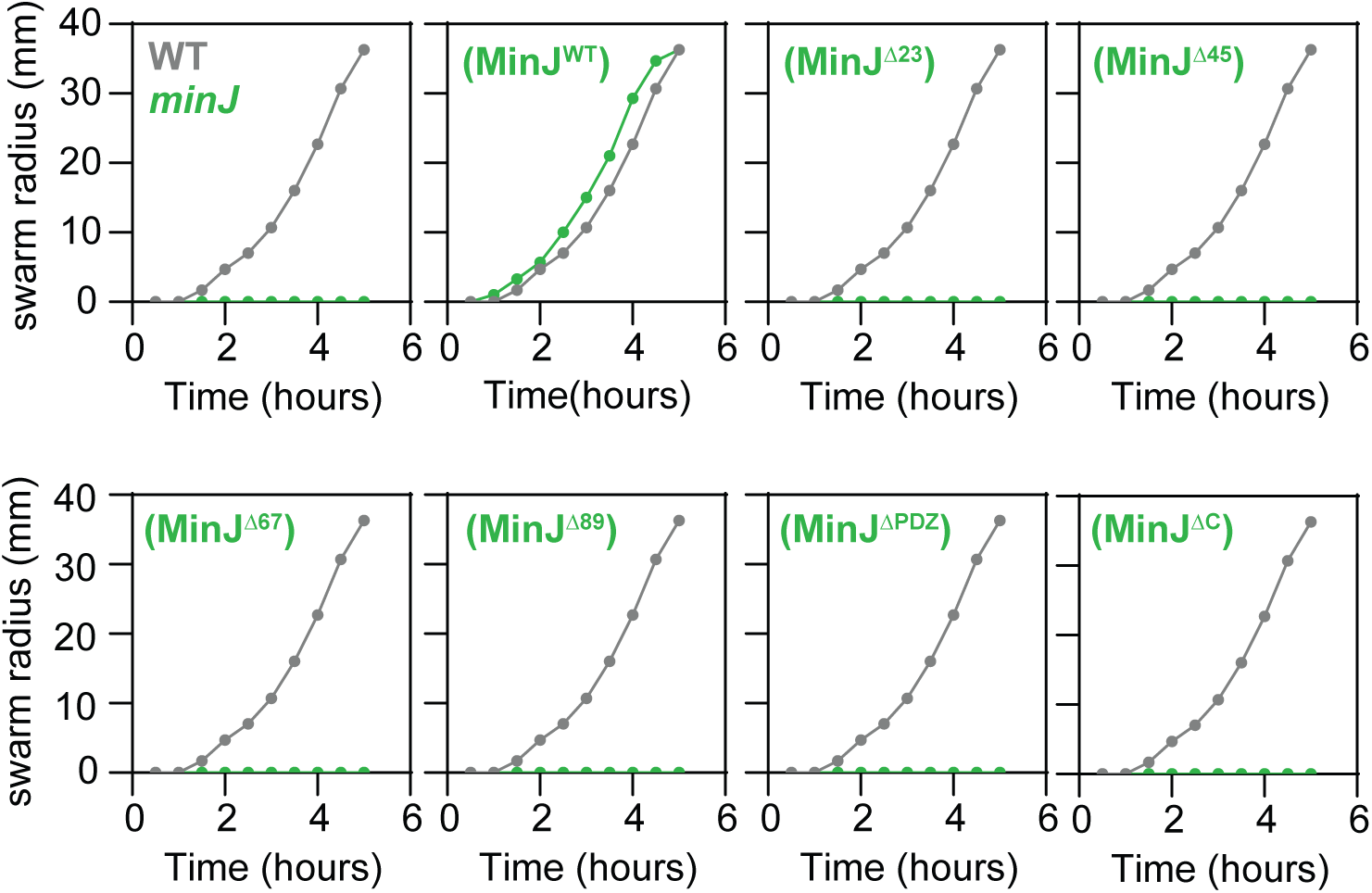
Each MinJ transmembrane domain and the C-terminal domain are required for swarming motility. Quantitative swarm expansion assays for the strains indicated in the upper right corner of each panel. The gray circle represents the wild type strain, while green circles represent complementation constructs expressed from the native minJ promoter and integrated at an ectopic chromosomal locus in a minJ mutant background. Whereas complementation with the wild type minJ allele restored swarming motility to the minJ mutant, complementation with the MinJ^Δ23^, MinJ^Δ45^, MinJ^Δ67^, MinJ^Δ89^, MinJ^ΔPDZ^, and MinJ^ΔC^ alleles failed to restore swarm expansion. The following strains were used to generate these data: WT (DK1042), *minJ* (DB92), *minJ (P_minJ_-minJ)* (DB2484), *minJ (P_minJ_-minJ^Δ23^)* (DB2499), *minJ (P_minJ_-minJ^Δ45^)* (DB2500), *minJ (P_minJ_-minJ^Δ67^)* (DB3144), *minJ (P_minJ_-minJ^Δ89^)* (DB2501), *minJ (P_minJ_-minJ^ΔPDZ^)* (DB2502), and *minJ (P_minJ_-minJ^ΔC^)* (DB2485).

**Figure S3.**
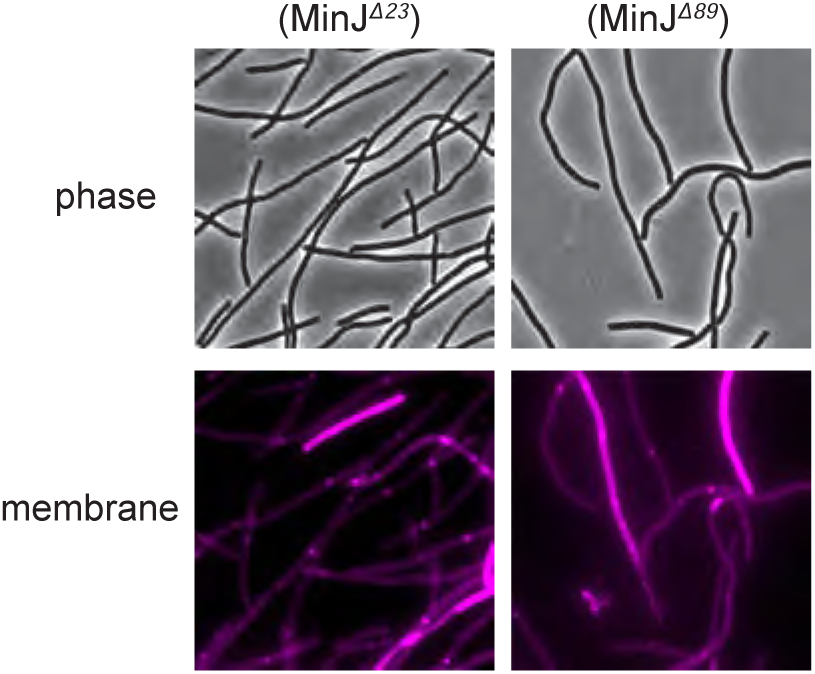
Deletion of MinJ transmembrane segments 2 and 3, or transmembrane segments 8 and 9, results in a cell division defect. Micrograph images of phase-contrast and fluorescence microscopy of cell membranes stained with FM4-64 (false colored magenta) for the genetic backgrounds indicated. The minJ mutant was complemented with either the *MinJ^Δ23^* or *MinJ^Δ89^* allele expressed from the native minJ promoter and integrated at an ectopic chromosomal locus. Both truncation alleles failed to complement the filamentous phenotype associated with loss of MinJ. The following strains were used to generate these data: *minJ (P_minJ_-minJ^Δ23^)* (DB2499), *minJ (P_minJ_-minJ^Δ89^)* (DB2501)

**Figure S4.**
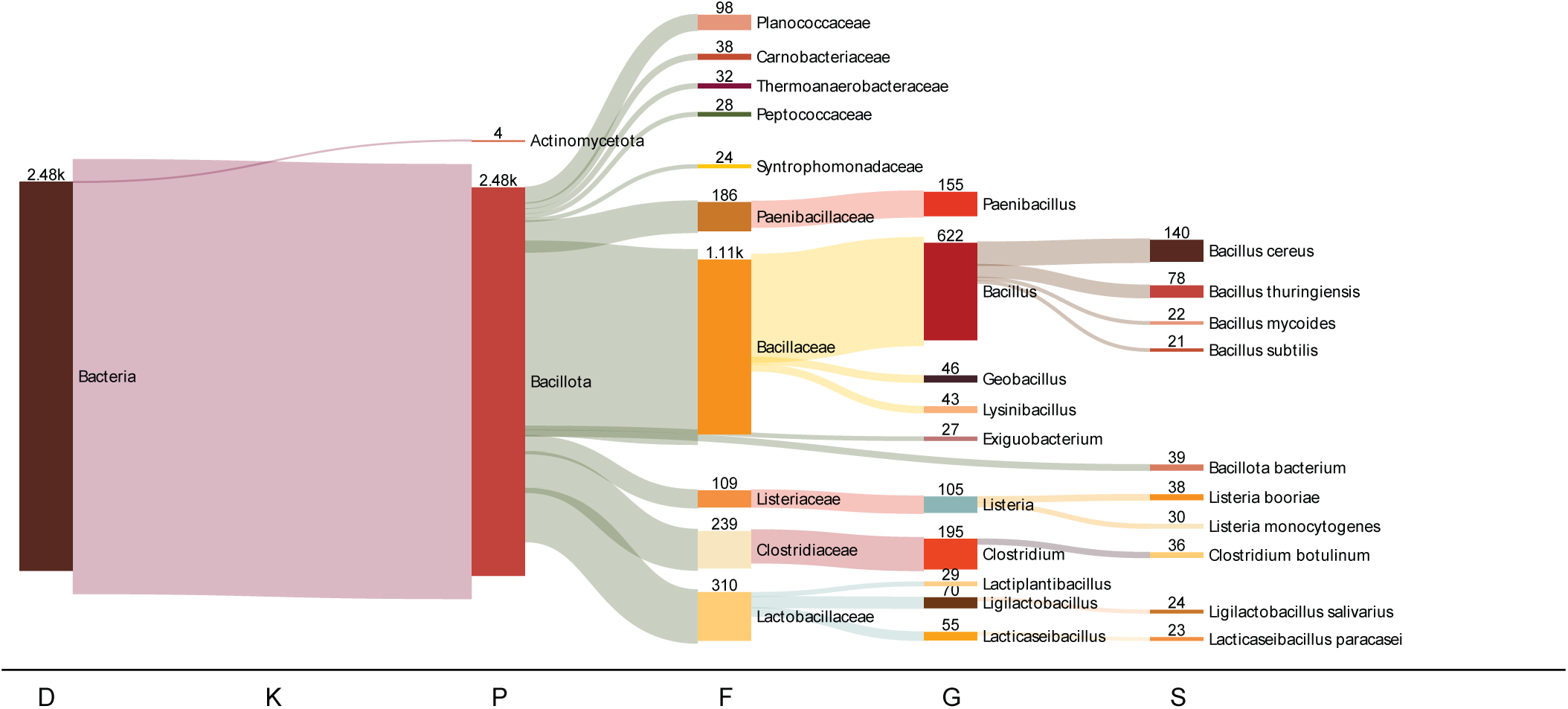
Foldseek structural clustering supports restriction of MinJ to the Bacillota. Foldseek cluster generated using the AlphaFold-predicted structure of *Bacillus subtilis* MinJ (UniProt accession O34775) as the query structure. The hierarchical clustering summarizes the taxonomic distribution of structurally related proteins recovered by Foldseek. Labels beneath each cluster indicate the number of sequences assigned to the corresponding taxonomic group. The cluster was overwhelmingly composed of proteins from the Bacillota, with 2,480 structural homologs identified within the phylum and only four distant matches assigned to the Actinomycetota. Progressive clustering is shown at the taxonomic levels of Domain (D), Kingdom (K), Phylum (P), Family (F), Genus (G), and Species (S).

**Figure S5.**
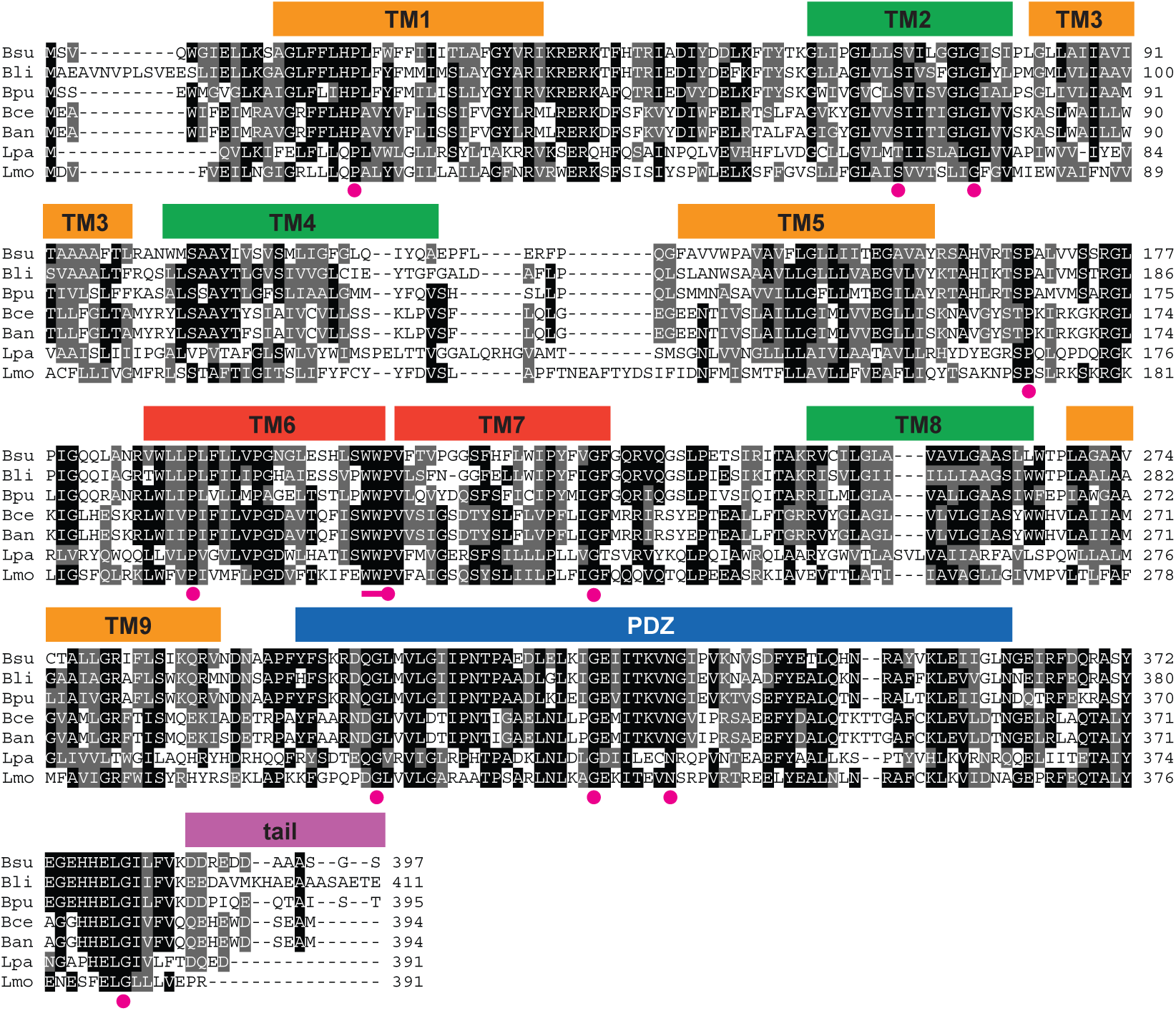
Conservation of MinJ topology and domain architecture across phylogenetically diverse orthologs. BoxShade multiple sequence alignment of representative MinJ orthologs from *Bacillus subtilis* (Bsu), *Lysinibacillus* (Bli), *Paenibacillus* (Bpu), *Bacillus cereus* (Bce), *Bacillus anthracis* (Ban), *Lactiplantibacillus plantarum* (Lpa), and *Listeria monocytogenes* (Lmo). Predicted transmembrane segments (TM1–TM9), the PDZ domain, and the C-terminal tail are indicated above the alignment. Golden-yellow boxes indicate transmembrane segments flanking cytoplasmic loops, green boxes indicate transmembrane segments flanking extracellular loops, and orange boxes indicate the predicted β-sheet transmembrane segments (TM6 and TM7). The PDZ domain is indicated in blue and the C-terminal tail in light purple. Conserved residues exhibiting greater than 90% conservation across the 396 MinJ orthologs are indicated by magenta circles. The highly conserved WWP motif (residues 207–209), exhibiting an average conservation greater than 80%, is indicated by the magenta arrow.

**Table S1.** Conserved residues across MinJ sequence.

| Residue Position | Residue | Percent Conserved (%) |
| --- | --- | --- |
| 20 | P | 90.68 |
| 70 | S | 93.20 |
| 77 | G | 91.44 |
| 168 | P | 97.48 |
| 191 | P | 96.73 |
| 209 | P | 91.94 |
| 228 | G | 96.22 |
| 305 | G | 96.47 |
| 325 | G | 96.47 |
| 332 | N | 96.22 |
| 380 | G | 98.99 |

<sup>a</sup>Residues with greater than 90% conservation identity

**Table S2.** Conserved sequence stretch across MinJ sequence.

| Residue position | Length | Consensus | Conservation |
| --- | --- | --- | --- |
| 207-209 | 3 | WWP | 81.11% |
| 206-209 | 4 | PWWP | 73.43% |
| 353-357 | 5 | CKLEV | 59.80% |

<sup>a</sup>Sequence stretches of three or more with greater than 50% conservation identity

**Table S3.**
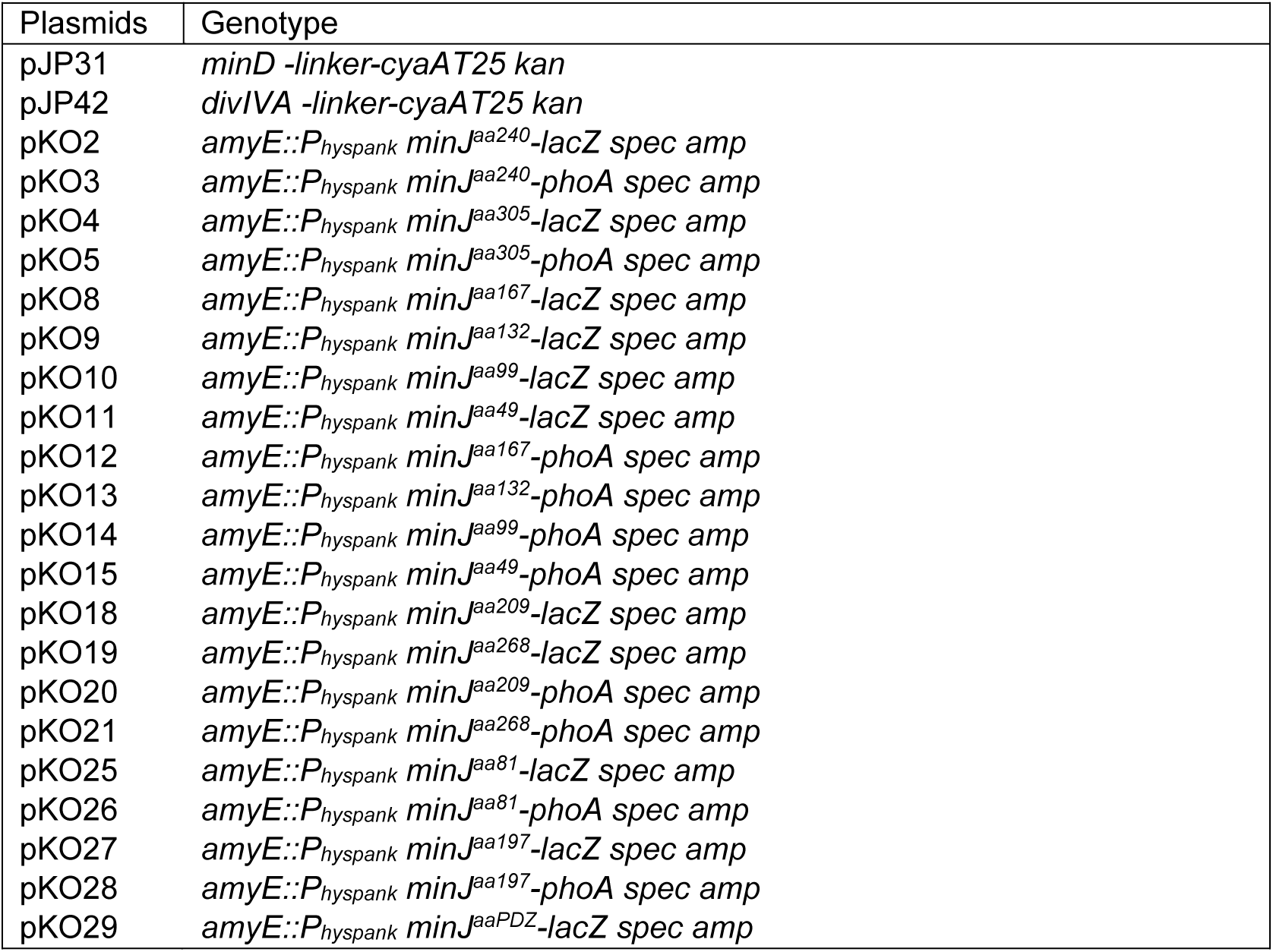
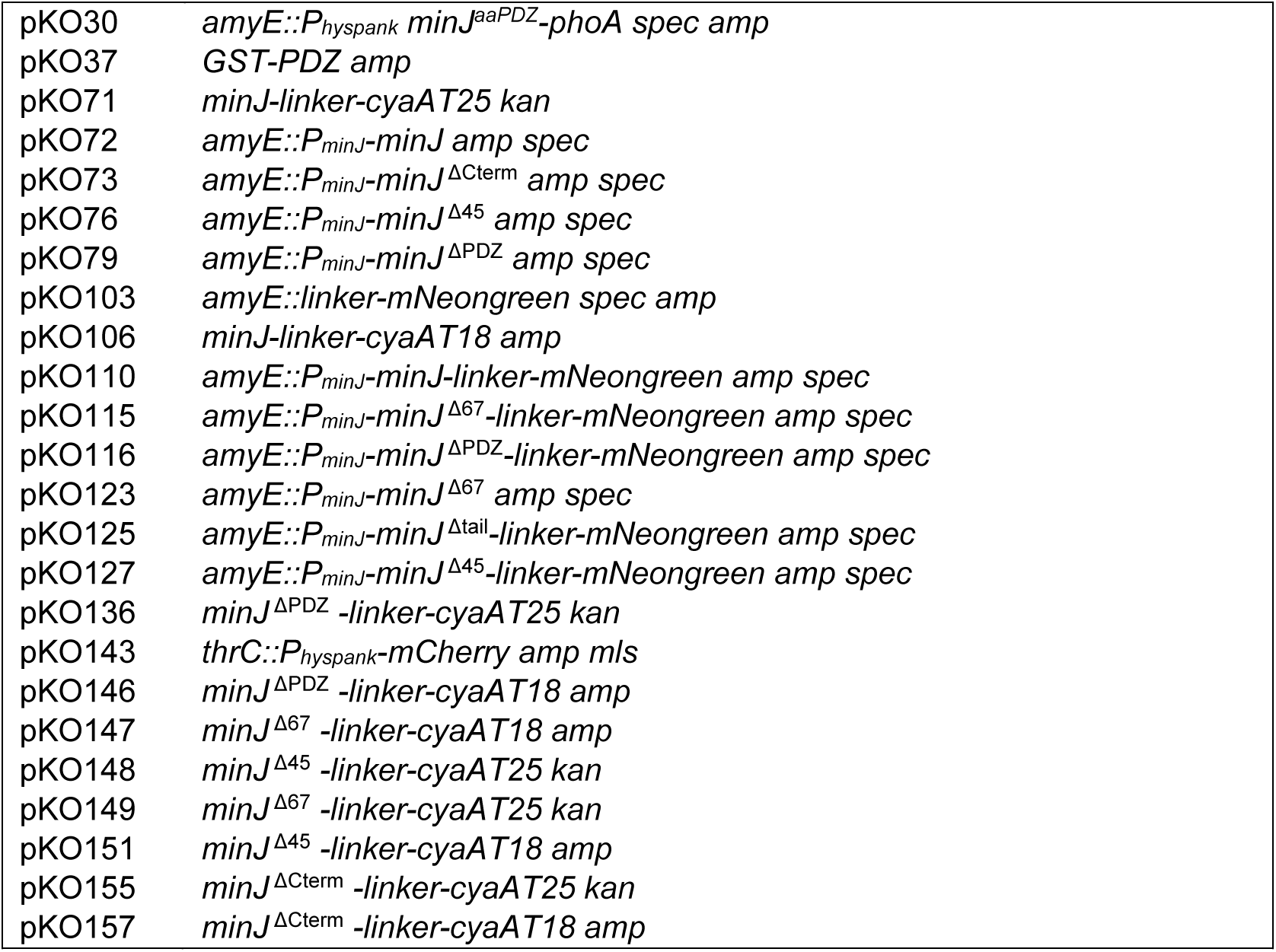
Plasmids.

**Table S4.**
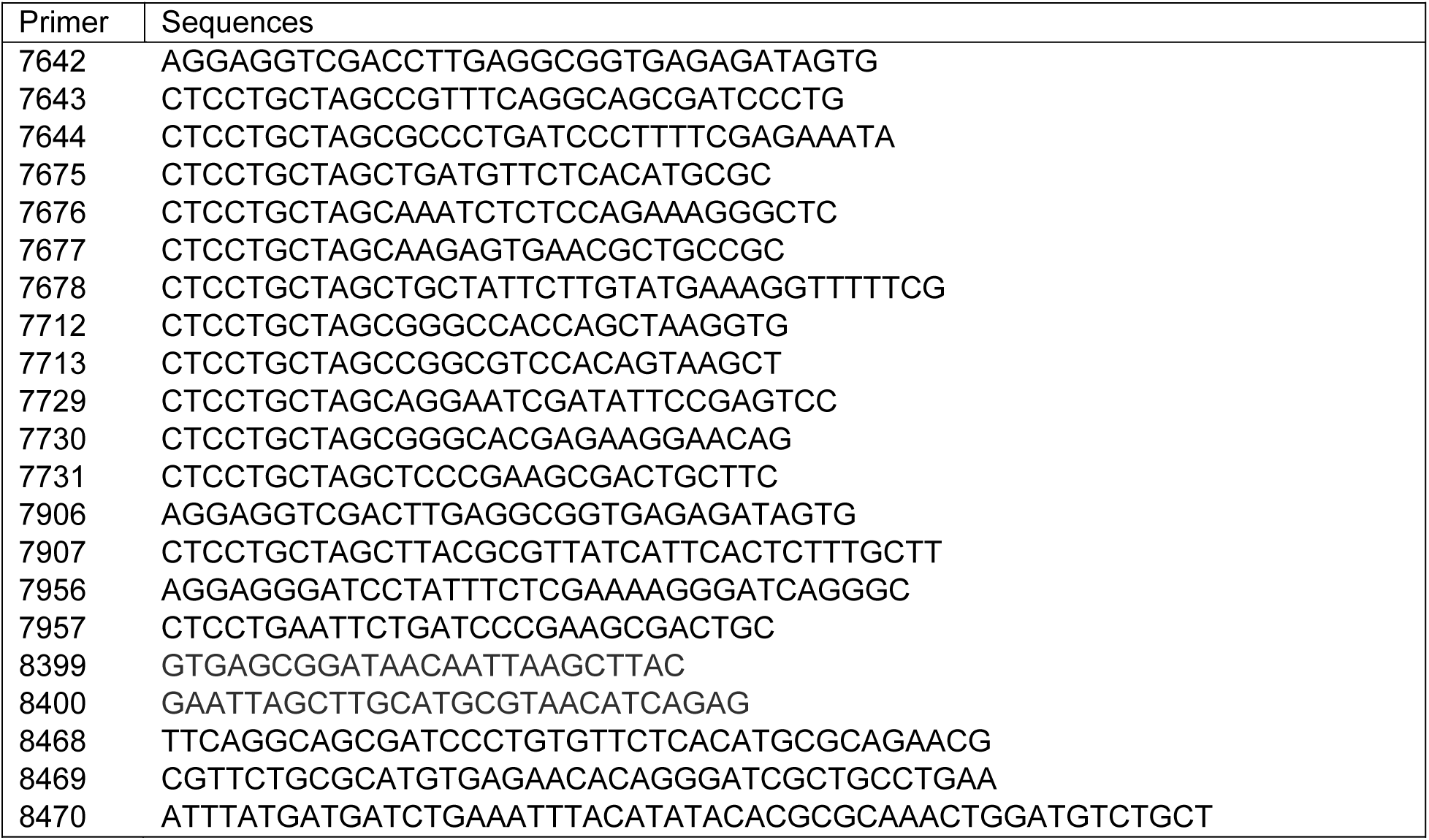
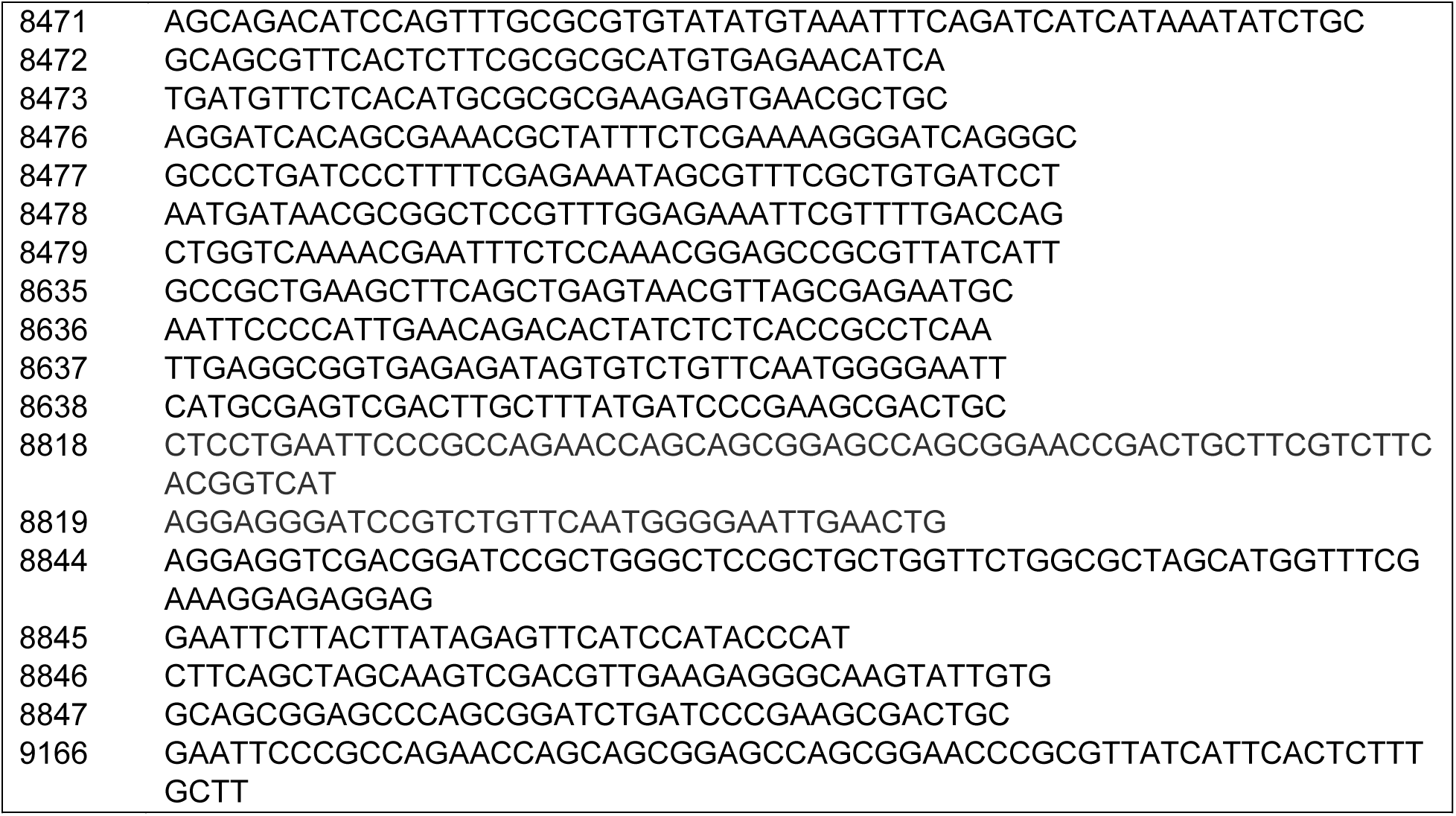
Primers.

